# Systematic Discovery of Pseudomonad Genetic Factors Involved in Sensitivity to Tailocins

**DOI:** 10.1101/2020.05.27.113977

**Authors:** Sean Carim, Ashley L. Azadeh, Alexey E. Kazakov, Morgan N. Price, Peter J. Walian, Romy Chakraborty, Adam M. Deutschbauer, Vivek K. Mutalik, Adam P. Arkin

## Abstract

Tailocins are bactericidal protein complexes produced by a wide variety of bacteria to compete against closely related strains. Like tailed bacteriophages, with whom they share an evolutionary and morphological relationship, tailocins bind and kill a narrow spectrum of target cells. Thanks to their high specificity, tailocins have garnered recent attention for their potential as precision antibacterial agents. Nevertheless, the field currently lacks a systematic investigation of genetic determinants of tailocin sensitivity. Here, we employed barcoded transposon-insertion mutant libraries and comparative genomics to assess genetic contributions to tailocin sensitivity in pseudomonads. Our mutant screens identified O-specific antigen (OSA) composition and display as most important in defining sensitivity to our tailocins. Additionally, the screens suggest lipopolysaccharide (LPS) thinning as a mechanism by which resistant strains can become more sensitive to tailocins. Our comparative genomics analyses show a loose relationship between OSA biosynthetic genes and tailocin sensitivity, as well as sensitivity nuances that require further investigation. Overall, our data reinforces the model that LPS molecules can act as either a receptor for, or shield against, tailocin binding and killing. This work offers insight into the specificity of tailocins and tailocin-mediated competition, informing the potential use of tailocins in microbiome manipulation and antibacterial therapy.

## Introduction

Interference competition between closely-related taxa is often mediated by the production and release of bacteriocins^1^. Bacteriocins are genetically encoded, ribosomally synthesized toxins that typically display a narrow killing spectrum^2^. The largest bacteriocins (2-10 MDa) are referred to as phage-tail like bacteriocins or tailocins, and these are evolutionarily and morphologically related to bacteriophage tails, type VI secretion systems (T6SS) and extracellular contractile injection systems (eCIS). They are encoded by single, contiguous biosynthetic gene clusters that resemble sequenced prophages^3^, but lack capsid, integrase and terminase genes. Tailocins are either R-type or F-type, depending on whether they resemble *Myoviridae* or *Siphoviridae* phages, respectively.

In the laboratory, tailocin production can be induced by applying DNA damaging agents^3^. After tailocin particles are assembled in the cytoplasm, they are released by autolysis of the producing cell through activation of a dedicated lysis cassette. Tailocin target specificity is defined by its receptor-binding proteins (RBPs): tail fibers, tail spikes and tail tips^4^. Recent studies have indicated that the RBPs are modular: they naturally undergo localized recombination^5^, and they can be swapped in and out manually to engineer target specificity^6–9^. Following recognition and binding, tailocins kill target cells with very high potency, with one to a few particles sufficient for killing a sensitive cell^6,10–12^. The mechanism of lethality is membrane depolarization, following insertion and puncture by the phage tail-derived structure^3^. Additionally, there are indications that tailocin production can mediate changes in bacterial community structure and diversity^13–15^. Thus they may be useful as a tool for targeted manipulation of microbiomes, with applications in both research and biotechnology.

Despite their importance and potential, our knowledge of cellular elements with which tailocins interact remains limited, an issue exacerbated by the scarcity of genetic tools for undomesticated bacterial isolates. Biochemical^16–18^, and forward genetics studies^7,8,13,19–21^ in model strains suggest that tailocins bind to specific lipopolysaccharide (LPS) moieties, and that loss of these results in resistance. Nevertheless, it remains unclear whether LPS composition is the only determinant of tailocin sensitivity, and whether this differs between tailocin types. Meanwhile, tailocin producing strains are resistant to their own tailocins (i.e., they avoid self-intoxication), but no systematic investigation for genetic factors responsible for this phenotype has yet been conducted. We aim to address the above knowledge gaps by leveraging resources developed previously by us and collaborators: 1) barcoded genome-wide transposon-insertion mutant libraries (RB-TnSeq)^22,23^ in environmental pseudomonads; and 2) a 130-member genome sequenced pseudomonad isolate library.

In this work, we first investigate tailocin-mediated killing among a set of 12 *Pseudomonas* isolates. We identify the tailocin biosynthetic clusters in these strains, then induce and partially purify their tailocins. We characterize a subset of these tailocin samples via proteomics and transmission electron microscopy (TEM), then use them to screen RB-TnSeq^22^ mutant libraries for differential tailocin sensitivity phenotypes in 6 separate tailocin-strain interactions. Our fitness assays: 1) establish O-specific antigen (OSA) composition and display as major determinants of tailocin sensitivity, and 2) show that disruptions to phospholipid retrograde and LPS anterograde transport weaken resistance to tailocin self-intoxication. Then, we examine the relationship between OSA biosynthetic genes and tailocin sensitivity at a larger scale, profiling sensitivity of our characterized tailocins across 130 genome sequenced pseudomonads. We find that strains with the same overall OSA cluster typically display the same tailocin sensitivity pattern. However, OSA gene content alone is unable to explain all observed variance in sensitivity. Overall, this work represents the first systematic effort to identify genetic factors involved in sensitivity to tailocins. Our findings offer a more comprehensive and nuanced view on how bacteria can alter their sensitivity to tailocin-mediated interference.

## Results

### Identification, induction and partial purification of tailocins

We first examined tailocin production and sensitivity in 12 *Pseudomonas* strains isolated from groundwater. These strains are classified into 10 species under the genus “Pseudomonas_E” according to the Genome Taxonomy Database (GTDB)^24^ (see <u>Table S1</u> for all strain information). To aid discussion, we will describe these 12 strains using codenames (‘Pse##’) defined in <u>Table S1</u>. We computationally searched for tailocin biosynthetic clusters (<u>Methods</u>) in the 12 strains and found them in all except Pse13. All clusters were inserted only between the genes *mutS* and *cinA*, a known hotspot for prophage integration^25,26^ (<u>Fig. S1</u>). These tailocin biosynthetic clusters vary in size (13.4 to 57.0kb) and appear to encode 1 to 4 tailocin particles of both R- and F-type (<u>Fig. S1</u>, <u>Fig. 1A)</u>. The R-type particles can be classified into subtypes Rp2, Rp3 or Rp4 based on their evolutionary relationships to different *Myoviridae* phages (<u>Fig. 1B</u>)^27^. Only Pse11 also encodes an F-type particle, and it resembles both the F2 pyocin of *P. aeruginosa* PAO1^28^ and the tail portion of *P. protegens* Pf-5 lambdoid prophage 06^29^.

**Figure 1.**
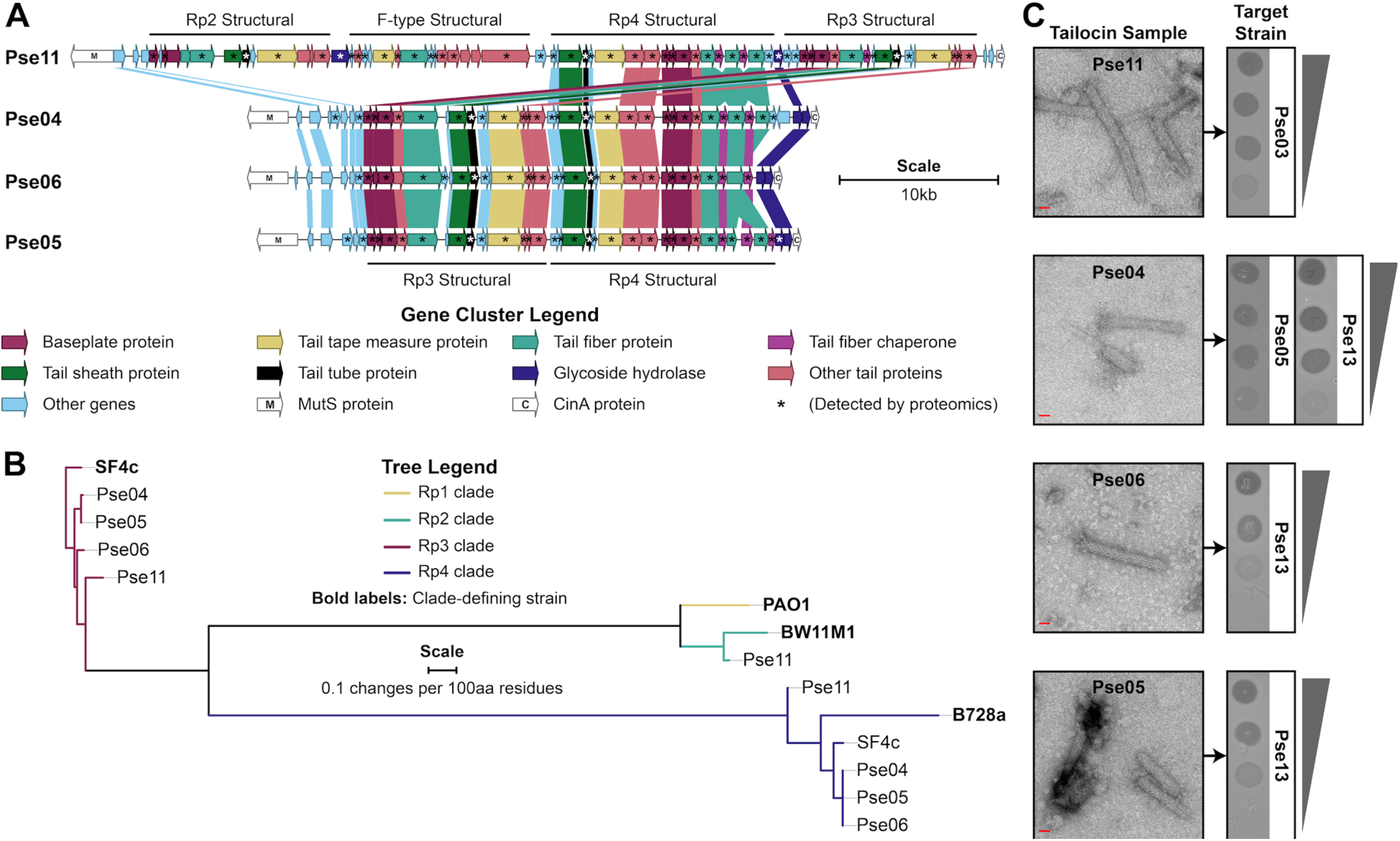
Characterization of select tailocin samples. Tailocins produced by our isolates Pse11, Pse04, Pse06 and Pse05 were characterized further in this study. **(A)** Tailocin biosynthetic gene clusters. Genes are colored by the presence of key words in their PHASTER^30^ predicted annotations. Genes in the same group of orthologs (“orthogroup”, see <u>Methods</u>) are joined by a block of color. The products of starred (*) genes are observed in our partially purified tailocin samples using MudPIT proteomics (<u>Methods</u>, <u>Table S7</u>). Additional data on these clusters can be found in <u>Table S4</u>. **(B)** Phylogeny of R-type tailocins by tail tube protein sequence. These are delineated by clade as per their relationship to the following tailocins: *P. aeruginosa* PAO1 R2 pyocin (Rp1), *P. putida* BW11M1 R-type tailocin (Rp2), *P. fluorescens* SF4c R-type tailocin 1 (Rp3) and *P. syringae* B728a R-type tailocin (Rp4)^27^. **(C)** Characterization of 5 lethal tailocin interactions by transmission electron microscopy and by spotting 5-fold dilution series of samples. Scale bar: 20nm. Grey triangles illustrate the relative concentration of tailocin in the dilution series. Zones of inhibition formed by diluted samples are indicative of non-replicative toxins, not phages. Spotting was done in triplicate and replicates gave identical data.

To assess whether these gene clusters are functional and responsible for encoding tailocins, we employed published protocols to induce and produce tailocins from our 12 strains and evaluate their targeting spectra^26^. Briefly, we induced tailocin production in each strain by applying mitomycin C to mid-log cultures and collected all tailocins from the culture supernatant through ammonium sulfate precipitation (<u>Methods</u>). As a control, we also subjected mitomycin C untreated culture supernatant to the same precipitation protocols. We then assessed the targeting spectra of these tailocin samples by spotting each sample on each of the 12 strains for a 12×12 killing matrix (<u>Fig. S2</u>). Consistent with the narrow target specificity expected of tailocins, lethality was sparse (15 killing interactions out of 144). Furthermore, no tailocin sample was toxic to its producing strain.

### Characterization of tailocin particles

We further characterized four of the tailocin samples (produced by Pse11, Pse04, Pse06 and Pse05). To examine the killing activity of our tailocins at lower concentrations, we spotted serial dilutions of the samples on sensitive strains. Spots made by the more diluted samples displayed reduced bacterial killing activity across the entire area of the zone of inhibition, and showed no distinct plaques (<u>Fig. 1C</u>). As reported earlier, this pattern is more typical of a diluted toxin, and not of a replicable killing agent like a phage^26^. To assess the protein composition of the tailocins and link them to structural genes, we subjected the samples to MudPIT LC/MS/MS proteomics analysis. Our proteomics data confirmed the presence of proteins for all possible tailocin particles encoded by our strains (<u>Fig. 1A</u>, <u>Table S7</u>). Thus, each sample likely consists of a mixture of 2 or 4 particles (<u>Fig. 1A</u>). Finally, we imaged each tailocin sample using transmission electron microscopy (<u>Fig. 1C</u>). Our micrographs showed an abundance of 20-25nm wide, straight or slightly curved rods, resembling past microscopy reports of *Pseudomonas* R-tailocins^10,17,31^. Most particles were uncontracted, with lengths between approx. 110 and 170nm. A few particles were contracted (see <u>Fig. 1C</u> Pse04 micrograph), with the 20-25nm wide segment (likely the tail sheath) visibly shortened, and with a narrower ∼18nm segment (likely the tail tube) protruding from one side.

To discern which tailocin particles are responsible for our observed killing activity, we aimed to delete the baseplate genes (<u>Fig. 1A</u>) for each of the R-type particles encoded by the four producing strains. Single baseplate locus deletion is a well-established method for eliminating assembly of a single tailocin particle while preserving the assembly of other particles produced by the strain^32,33^. We succeeded in generating 7 of the 9 possible baseplate mutants, partially purified tailocin samples from each mutant, and spotted them on sensitive strains (<u>Methods</u>, <u>Fig. S3</u>). Killing activity was assigned to a particle by identifying which baseplate mutation eliminates killing achieved by the wild-type. We found that Pse05 Rp3 kills Pse13, Pse04 Rp3 kills Pse05, Pse04 Rp4 kills Pse13, and both Pse06 Rp3 and Rp4 kill Pse13. Additionally, we observe that killing of Pse03 by Pse11 is not mediated by the latter’s Rp4 particle, but are unable to specify which of its remaining particles are responsible (<u>Fig. S3</u>). In the case of Pse04, we corroborate findings that encoding multiple tailocin particles increases the range of susceptible targets^32^.

### Genome-wide mutant fitness assays identify gene functions involved in tailocin sensitivity

To study genes important in tailocin sensitivity, we sourced three pooled, barcoded, genome-wide transposon-insertion (RB-TnSeq) mutant libraries (for strains Pse05, Pse03 and Pse13) reported previously^23^. We then performed fitness experiments on these libraries, assaying five library and tailocin sample combinations (see <u>Fig. 1C</u> and <u>Fig. 2A</u>). In these experiments, selection pressure from the tailocins increases the relative abundance of resistant mutants. Changes in the relative abundance of mutants is monitored using next-generation sequencing via the BarSeq approach^22^ to track DNA barcodes before and after tailocin treatment. This barcode sequencing data is calculated into fitness scores for each strain and each gene as described earlier^22^ (<u>Fig. 2A</u>, <u>Methods</u>). Here, the positive gene fitness scores indicate genes conferring relative tailocin *resistance* when disrupted. Conversely, negative fitness scores indicate genes that confer relative tailocin *sensitivity* when disrupted.

**Figure 2.**
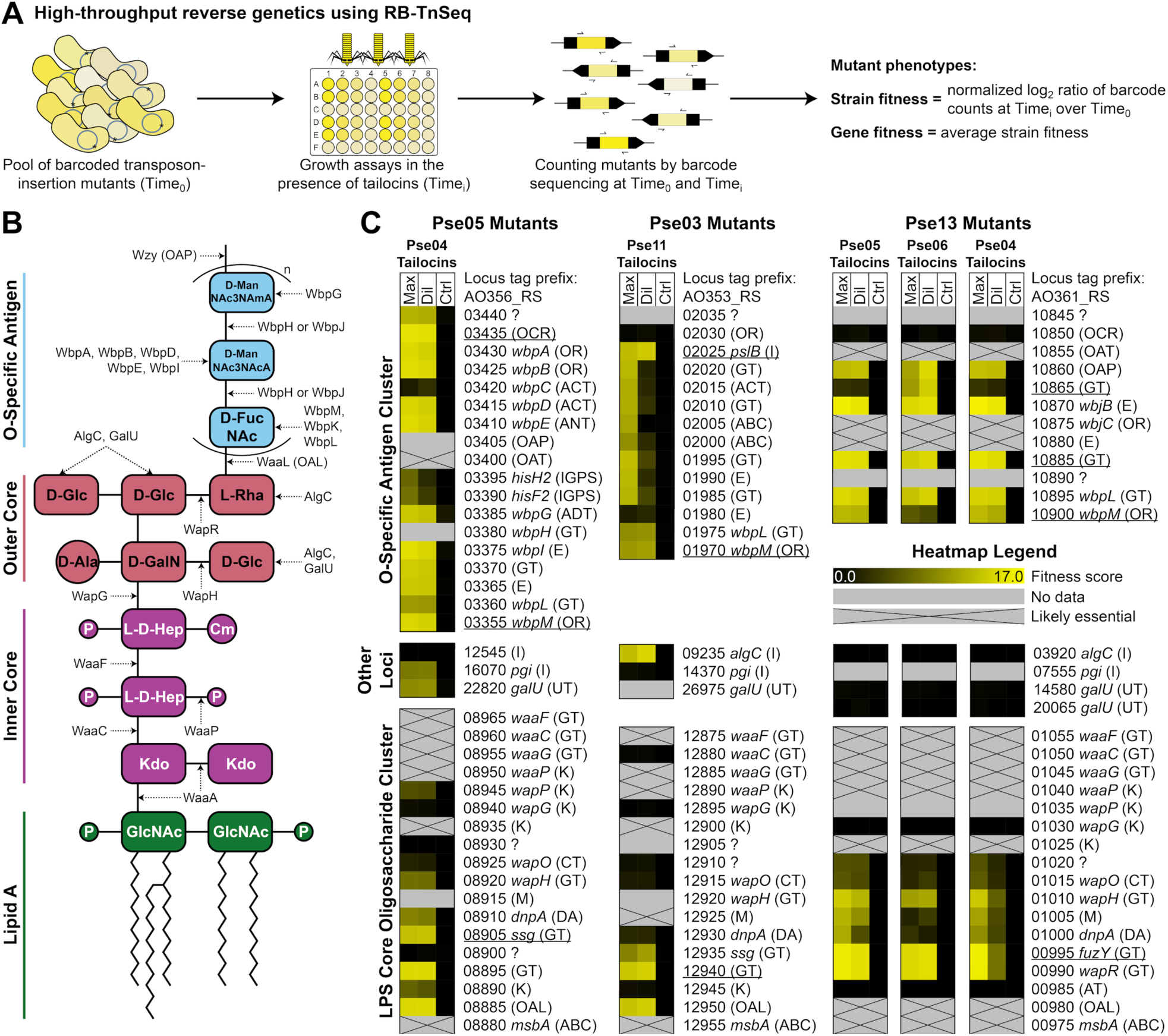
Genes implicated in tailocin sensitivity. **(A)** The RB-TnSeq approach for measuring normalized gene fitness. **(B)** Structure of the O-specific antigen LPS of *P. aeruginosa* PAO1 and enzymes involved in its assembly, based on King et al. (2009)^34^. Chemical moieties are abbreviated with the following definitions: Cm, O-carbamoyl; D-Ala, D-alanine; D-FucNAc, 2-acetamido-2-deoxy-D-fucose; D-GalN, D-galactosamine; D-Glc, D-glucose; D-ManNAc3NAcA, 2,3-diacetamido-2,3-dideoxy-D-mannuronate; D-ManNAc3NAmA, 2-acetamido-3-acetamidino-2,3-dideoxy-D-mannuronate; GlcNAc, N-acetyl-D-glucosamine; Kdo, ketodeoxyoctonoate; L-D-Hep, L-glycerol-D-manno-heptose; L-Rha, L-rhamnose; P, phosphorylation (with either one or multiple phosphates or phosphoethanolamines). Select gene function abbreviations are also included: OAL, O-antigen ligase; OAP, O-antigen polymerase. **(C)** Heatmaps depicting fitness scores for genes that confer tailocin resistance when mutated. Tailocins were supplied at two different concentrations: maximum concentration (‘Max’) and a 10-fold dilution from maximum (‘Dil’). Tailocin-free buffer was supplied for control experiments (‘Ctrl’). All fitness assays were performed in duplicate, and the average fitness is displayed. A gene lacking data means we did not obtain a transposon insertion in it^23^. Genes were annotated in two levels of detail depending on their homology to well-characterized genes. First, all genes were labeled with a function abbreviation. ABC, ABC transporter subunit; ACT, acetyltransferase; ADT, amidotransferase; ANT, aminotransferase; AT, acyltransferase; C, carbamoyltransferase; DA, deacetylase; E, epimerase; GT, glycosyltransferase; I, isomerase; IGPS, imidazole glycerol phosphate synthase subunit; K, kinase; M, Mig-14-like; OAL, O-antigen ligase; OAP, O-antigen polymerase; OAT, O-antigen translocase; OCR, O-antigen chain length regulator; OR, oxidoreductase; UT, uridylyltransferase; ?, unknown. Then, if a gene’s translation shared ≥60% identity with ≥90% coverage to a characterized gene in *P. aeruginosa* PAO1, *P. aeruginosa* PA103 or *P. fluorescens* SBW25, it was assigned the same gene name. Phenotypes of <u>underlined</u> genes have been validated by spotting tailocin samples on individual mutants (<u>Fig. S4</u>). See <u>Tables S9 to S17</u> for read count, t-like statistic and additional fitness data for these genes.

Treatment with an antagonizing tailocin results in a few very high, replicable, positive gene fitness scores (up to +16), indicative of reliable selection for a very small subset of mutants. These genes are clearly important for tailocin sensitivity. However, the competitive advantage of the high fitness mutants in our pooled assays means mutants with weaker positive fitness have likely been severely outcompeted and eliminated from the population. Our earlier work with phage fitness assay experiments^35,36^, which, like here, involve the application of a stringent selective agent, suggests we focus our discussion on gene disruptions that confer the top positive fitness phenotypes in the presence of tailocins (see <u>Methods</u> for criteria). Genes that passed our criteria were few in number (44 total genes in 3 strains across 5 fitness assays) (<u>Tables S10</u>, <u>S13</u> and <u>S16</u>). All but three of these genes lie within either the LPS core oligosaccharide biosynthetic gene cluster or the O-specific antigen (OSA) biosynthetic gene cluster. The three remaining genes (orthologs of *pgi, galU* and *algC*), while located elsewhere in the genome, are involved in the assembly of LPS monomers^34^. Notably, no gene disruptions in our strains’ common polysaccharide antigen (CPA) clusters (Pse05 AO356_RS11820-80; Pse03 AO353_RS10215-75; Pse13 AO361_RS11625-90) which are responsible for assembling a separate O-polysaccharide molecule^34^, gave high enough fitness scores to pass our criteria.

While there is considerable heterogeneity in LPS gene content across Pse05, Pse03 and Pse13, we can still make some inferences about their functions using homology to well-characterized PAO1 LPS genes. The LPS core gene clusters in our strains are generally homologous to PAO1 and to each other (<u>Fig. S5</u>), so those genes are mostly assigned high confidence annotations as a result (see <u>Fig. 2C</u> caption). Variation in the LPS core clusters include the absence of *wapP* in Pse03 and the weak homology of the *wapR* orthologs in Pse05 and Pse03. Meanwhile, only Pse05 has an OSA gene cluster homologous to PAO1 (<u>Fig. S6</u>), making that cluster the best annotated of its type on <u>Fig. 2C</u>.

### The O-specific antigen as the key factor in tailocin sensitivity

In Fig. 2C, we depict fitness score data for all genes in the LPS core and OSA clusters, and homologs of *algC, pgi* and *galU*. Thus, <u>Fig. 2C</u> allows comparison of LPS biosynthetic genes by their importance in tailocin-mediated killing. Overall, the high fitness genes encode proteins that: build LPS core monomers (*galU, algC*), assemble the LPS core (*wapH, wapR*), build OSA monomers (*wbpA, wbpB, wbpD, wbpE, wbpG, wbpI, wbpL, wbpM, pgi, pslB, algC, wbjB*), polymerize OSAs, regulate the length of OSAs, translocate OSAs, and ligate OSAs to the LPS core. Loss of any of these functions in *P. aeruginosa* is known to result in truncation or complete absence of one or both of the OSA and CPA^34^, or, in the case of the chain length regulator, display of OSAs with altered length^34^. This suggests that, in our strains, intact, correct length OSA is necessary for tailocin-mediated killing, likely as a receptor for tailocin binding. This concurs with, and expands on, recent work with Rp4 tailocins in *P. syringae*^*21,37*^.

Since there are high fitness disruptions in orthologs of genes involved directly in LPS outer core assembly (*galU, wapH, wapR*), one could argue that some LPS outer core residues also serve as tailocin receptors. While there is evidence that other *Pseudomonas* tailocins target the outer core^13,18,38^, the strong benefits of OSA mutations imply that OSA is the primary receptor for our studied tailocins. The phenotypes of (*galU, wapH, wapR*) disruption could reflect the loss of OSA, as the structures they assemble are required for OSA attachment^34^. Thus we only make claims about the role of the OSA in this work. For discussion of the role of LPS inner core biosynthetic enzymes, see <u>Supplementary Notes</u>.

Phages are known to use both LPS moieties and membrane proteins as receptors^39^, but tailocins are only known to use LPS moieties as receptors^13,16,18,20,21,27,37,38^. Our data, which represents the most comprehensive genetic analysis so far of possible tailocin receptors, maintains this paradigm. In work recently published by ours and other groups^35^, RB-TnSeq and a similar fitness assay design is used to find genes required for sensitivity to *Escherichia coli, Phaeobacter inhibens* and *Salmonella enterica* phages^35,36,40^, and the data recovers their known membrane protein receptors. Thus, our methods employed here are capable of identifying non-essential membrane protein receptors, and the tailocins we tested do not depend on such factors.

### Disruption of specific OSA biosynthesis genes confers resistance to a subset of antagonistic tailocins

Challenging the Pse13 RB-TnSeq library with three different antagonistic tailocin samples allowed us to identify genes that are involved in sensitivity to some, but not all, of these tailocins. We identified two such genes, AO361_RS10865 and AO361_RS10900, both in the OSA biosynthetic cluster (<u>Fig. 2C</u>). Disruption of AO361_RS10865, which is a putative glycosyltransferase, confers greatly increased resistance to Pse06 tailocins, but has a small effect on resistance to Pse05 and Pse04 tailocins. In contrast, disruption of AO361_RS10900, an ortholog of WbpM (UDP-4,6-GlcNAc dehydratase) in *P. aeruginosa* PAO1 (78% identity, 100% coverage), confers resistance to Pse05 and Pse04 tailocins, but has a reduced effect on killing by Pse06 tailocins (<u>Fig. 2C</u>). We validated these phenotypes by spot tests on individual Pse13 mutants (<u>Fig. S4</u>). See <u>Supplementary Notes</u> for further review of the putative functions of AO361_RS10865 and AO361_RS10900. Our findings highlight the speed and fine resolution of RB-TnSeq fitness assays in identifying genes involved in tailocin resistance in non-model bacteria.

### Disruption of genes encoding outer membrane lipid asymmetry and LPS transport functions increases sensitivity to tailocin self-intoxication

A wide body of evidence shows that tailocin-producing strains are resistant to the tailocins they produce^5,13^, in a phenotype referred to as resistance to self-intoxication (<u>Fig. 3A</u>, <u>Fig. S2</u>). Tailocins are believed to have evolved from prophages, so resistance to self-intoxication may have arisen from natural selection away from self-targeting, and toward targeting competing bacteria. Mechanistically this is thought to occur through evolution of the tailocin’s receptor-binding proteins (RBPs)^3^. Unlike with S-type bacteriocins, which are genetically transferred along with a cognate immunity gene, there are no known immunity genes for tailocins.

**Figure 3.**
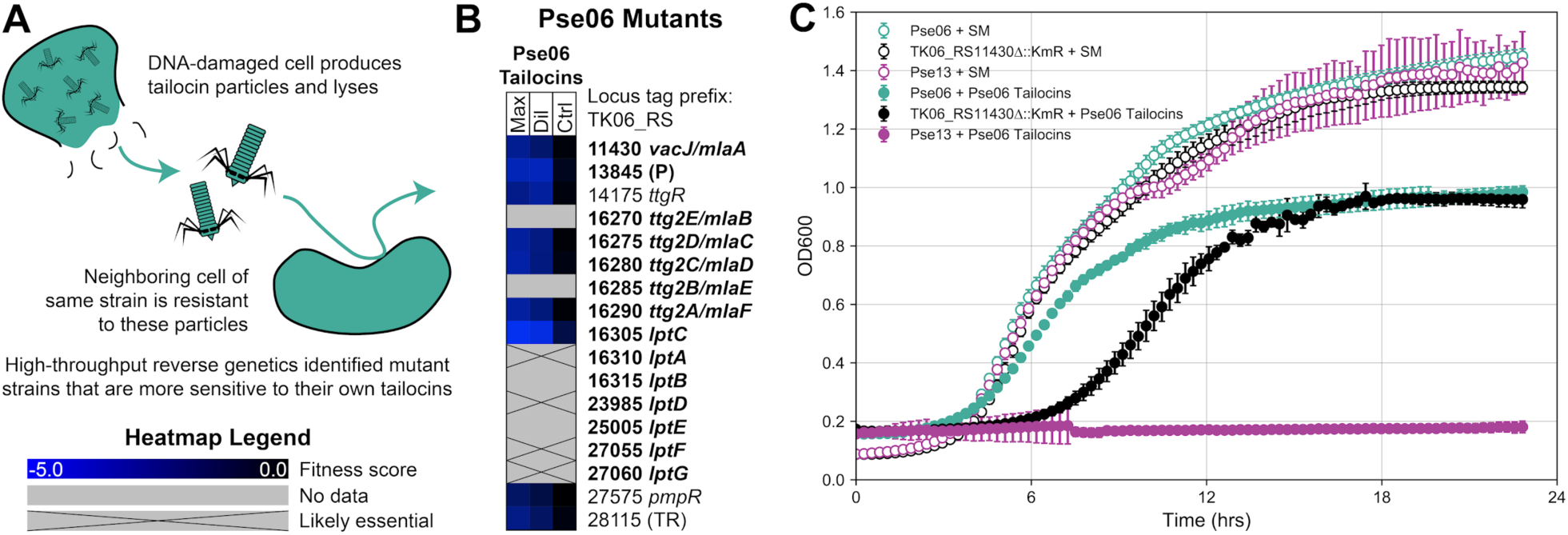
Genes implicated in resistance to tailocin self-intoxication. **(A)** Illustration of resistance to tailocin self-intoxication. **(B)** Heatmap depicting fitness data for transposon-insertion mutations in Pse06 that confer negative fitness when challenged by Pse06 tailocins. Tailocins were supplied at two different concentrations: maximum concentration (‘Max’) and a 10-fold dilution from maximum (‘Dil’). Additionally, tailocin-free buffer was supplied for control experiments (‘Ctrl’). A gene lacking data means we did not obtain a transposon insertion in it^23^. Genes were annotated with gene names from either *P. aeruginosa* PAO1 or *P. putida* KT2440 if their amino acid sequence shared ≥60% identity with ≥90% coverage. *lptC* was additionally labeled thanks to its genomic context. Remaining genes were annotated with a function prediction: P, peptidase; TR, transcriptional repressor. **Bold** genes are putatively involved in maintaining LPS display at the outer leaflet of the outer membrane. See <u>Tables S18 to S20</u> for read counts, t-like statistics and additional fitness data for these genes. **(C)** Phenotypic validation by growth curve for the independently generated TK06_RS11430 (*vacJ/mlaA*) deletion in Pse06. SM, SM buffer. The mutant displays increased sensitivity to Pse06 tailocins compared to Pse06 wild-type. Pse13 growth curves are included as a tailocin-sensitive control. These experiments were repeated (triplicate) and averages are plotted. Error bars: standard deviation.

To uncover genes involved in resistance to tailocin self-intoxication, we challenged the Pse06 RB-TnSeq library with tailocins produced by wild-type Pse06, and looked for strong negative fitness (≤-2.0) effects. That is, we looked for mutants that show increased sensitivity when treated with tailocins as compared to the no-treatment control. These criteria yielded 9 hits, depicted in <u>Fig. 3B</u>. 6 of the 9 hits appear to have functions relevant to maintaining a sufficient density of LPS at the outer leaflet of the outer membrane (OM) (see bold genes in <u>Fig. 3B</u>)^41,42^. Among these, 4 of the hits (TK06_RS11430, TK06_RS16275, TK06_RS16280, TK06_RS16290) are in the *mla* (maintenance of lipid asymmetry) pathway, which constitute a retrograde phospholipid (PL) transport system involved in OM lipid homeostasis. The *mla* pathway removes PLs that mislocate to the outer leaflet of the OM so that dense packing of LPS, and OM integrity, is maintained^43^. Another hit is an ortholog of *lptC* (TK06_RS16305), encoding a component of the *lpt* lipopolysaccharide transport system that delivers LPS to the outer leaflet of the OM^44^. One more hit is a putative metalloprotease (TK06_RS13845) with weak homology (36% identity, 95% coverage) to *E. coli*’s BepA, a periplasmic chaperone or protease for outer membrane proteins. BepA assists in the formation of LptD, an OM beta-barrel protein, and also degrades misassembled LptD^45^. Finally, the remaining 3 sensitivity hits– TK06_RS14175 (repressor of multidrug efflux pump^46^), TK06_RS27575 (quorum sensing response regulator^47^) and TK06_RS28115 (TetR family transcriptional regulator)–do not have discernible functions in LPS display. None of our 9 hits localize to the tailocin biosynthetic cluster. This suggests that tailocins do not possess co-located immunity factors, which is common for other proteinaceous toxins (e.g. S-type bacteriocins, lactic acid bacteria type I and II bacteriocins, and type IV and VI secretion system effectors)^2,48–51^.

To validate these increased tailocin sensitivity phenotypes, the 9 inferred sensitizing mutations were independently generated (see <u>Methods</u>), and the tailocin sensitivity phenotypes of these mutants were validated by measuring growth curves of planktonic cultures in the presence and absence of Pse06 tailocins. An exemplar set of growth curves for the TK06_RS11430 mutant is illustrated in <u>Fig. 3C</u>. Additional growth curves for the remaining 8 disruptions are presented in <u>Fig. S7</u>. In each case, these mutants exhibited impaired growth relative to wild-type Pse06 in the presence of the Pse06 tailocins. However, the mutants do not become completely sensitive to Pse06 tailocins, like Pse13, a strain included as a sensitive control.

### The relationship between OSA biosynthetic gene content and tailocin sensitivity across an isolate library

Since our mutant fitness data points to OSA composition and display as the key factors in tailocin sensitivity, we decided to examine the relationship between a target strain’s OSA biosynthetic gene cluster and its tailocin sensitivity among natural strain variants. To do this, we expanded sensitivity phenotyping of our four tailocin samples (<u>Fig. 1C</u>) to our in-house collection of 128 genome sequenced groundwater *Pseudomonas* isolates (classified into 29 GTDB species under the genus “Pseudomonas_E”^24^, <u>Table S1</u>). We also included the well studied human pathogen *P. aeruginosa* PAO1 and plant pathogen *P. syringae* B728a in our sensitivity assays. All 130 strains were hierarchically clustered by their OSA gene content similarity (see <u>Methods</u>). <u>Fig. 4</u> shows the tailocin sensitivity data decorated onto the OSA clustering of the 130 strains. Separately, we arranged the strains onto a phylogenetic tree built from a concatenation of 88 single-copy marker gene translations to compare phylogenetic distance with tailocin sensitivity (<u>Fig. S8</u>, <u>Methods</u>).

**Figure 4.**
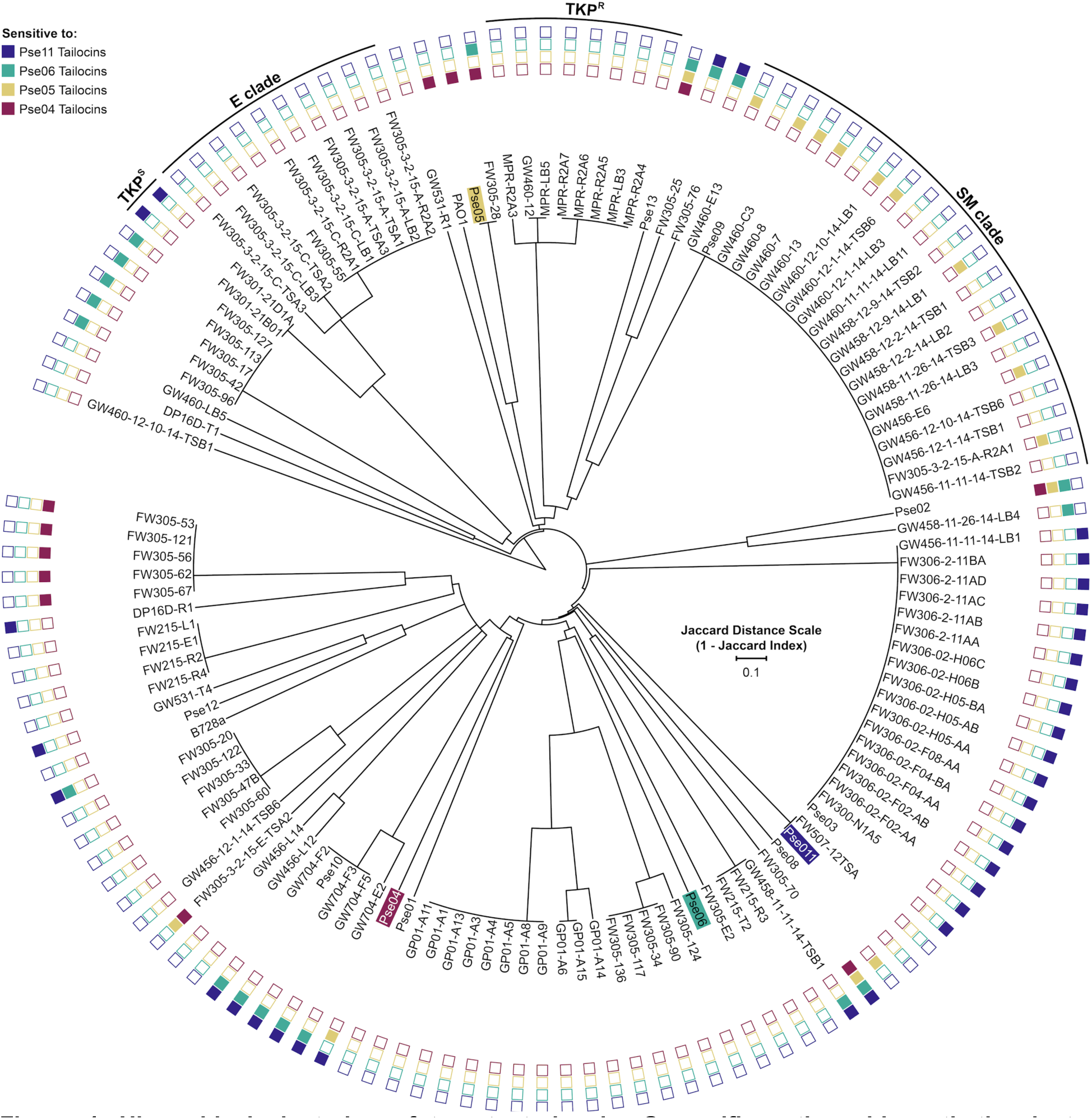
Hierarchical clustering of target strains by O-specific antigen biosynthetic cluster similarity, overlaid with tailocin sensitivity data. 130 *Pseudomonas* strains were clustered by the similarity of their OSA clusters, assessed as Jaccard distance of orthogroups (OrthoFinder v2.2.7^52^) found in the OSA cluster genes. See <u>Methods</u> for details on our approach to delineating OSA clusters. Tailocin producing strain labels are highlighted in color. Shaded boxes at the outer edge of the tree indicate sensitivity of that strain to the correspondingly colored tailocin. To aid discussion of 3 clades of strains with interesting features, we named those clades after the most closely related species by 16S rRNA similarity: E clade (*P. extremorientalis*); TKP clade (*P*. sp. TKP); SM clade (*P. silesiensis*/*P*.*mandelii*). In this clustering, TKP strains are separated into two groups: TKP^S^ (Pse11 Tailocins sensitive) and TKP^R^ (resistant). For a phylogenetic tree of these strains, see <u>Fig. S8</u>. For a comparative illustration of the OSA clusters encoded by these strains, see <u>Fig. S9</u>. For the killing matrix in table format, see <u>Table S6</u>.

Generally, strains with similar or identical OSA clusters had the same tailocin sensitivity phenotypes. This can be seen across <u>Fig. 4</u> or <u>Fig. S9</u> as members of the same dendrogram branch sharing the same pattern of sensitivity to the four tailocin samples. An exception to this is the ‘SM’ clade, whose 21 members shared identical (to the nucleotide level) OSA genes (<u>Fig. S9</u>) and a close phylogenetic relationship (>99.9% pairwise average nucleotide identity, ANI) but vary in their sensitivity to Pse05 tailocins (11 sensitive, 10 resistant). We compared these strains at their *mla* and *lpt* loci and found that those, too, are identical. Finally, we performed a genome-wide association study (GWAS) on the SM strains using DBGWAS^53^ which failed to find any genomic factors associated with Pse05 tailocin sensitivity with a q-value below 0.64. At this point, the genotypes responsible for the differential tailocin sensitivity in the SM clade are unknown to us, and additional work is needed to uncover them.

Additionally, strains that cluster by OSA similarity are also typically similar across most of the genome (<u>Fig. 4</u> and <u>Fig. S8</u>), though again there are exceptions. The most notable exception here is among members of the ‘TKP’ clade, all of whom group together phylogenetically (<u>Fig. S8</u>). 2 of the 10 TKP strains (TKP^S^) display sensitivity to Pse11 tailocins by spot test, while the remaining 8 strains (TKP^R^) are resistant. Comparison of the OSA clusters between TKP^S^ and TKP^R^ strains shows that they are highly divergent (<u>Fig. S9</u>). Curiously, the OSA clusters of TKP^S^ strains closely resemble those encoded by strains in the ‘E’ clade (which is resistant to Pse11 tailocins), but differ in one region that putatively encodes two hypothetical proteins and a sugar-modifying enzyme (<u>Fig. S9</u>). Since loss of a single OSA biosynthetic gene can alter the tailocin sensitivity of a strain (<u>Fig. 2C</u>), we hypothesize that the genes in this variable region are responsible for the differential sensitivity of TKP strains to Pse11 tailocins. The opposite scenario is exemplified by Pse05 and PAO1, which are phylogenetically distinct (79.95% ANI), yet share similar OSA clusters and the same tailocin sensitivity pattern (<u>Fig. 4</u>).

Our comparative genomics analysis is in incomplete agreement with our fitness data, which emphasizes the importance of OSA on tailocin sensitivity. A loose relationship between OSA and sensitivity can be seen: strains with the same overall OSA cluster tend to have the same sensitivity. However, we also observe that variation in sensitivity can be achieved without apparent change in OSA gene sequence, exemplified by the SM strains. Thus, factors involved in tailocin sensitivity are more complex and nuanced than what we have been able to uncover here, and future experiments must be designed with this in mind.

## Discussion

In this work, we first systematically analyze the tailocin encoding capabilities of *Pseudomonas* isolates, then partially purify and characterize the encoded tailocins. We then use our RB-TnSeq technology to uncover genetic determinants of tailocin sensitivity in 6 different tailocin-strain interactions. Our results highlight the role of O-specific antigen composition and display as the key factors defining tailocin sensitivity, and do so at gene-level resolution. We identify individual OSA biosynthetic genes whose functions are involved in sensitivity to different specific tailocins. Subsequently, we assemble evidence supporting the role of lipopolysaccharide density in tailocin resistance, showing that mutations to retrograde phospholipid transport and anterograde LPS transport systems increase the sensitivity of a strain to its own tailocins. Finally, we investigate the relationship between OSA gene content and tailocin sensitivity across 130 *Pseudomonas* strains. Strains with highly similar OSA clusters broadly share the same tailocin sensitivity pattern, even if they are phylogenetically distant, but exceptions exist. Our analysis suggests that genetic factors beyond those studied here are involved in tailocin sensitivity, warranting more nuanced investigation. We summarize the major genetics findings in this study by proposing five scenarios that illustrate the effect of OSA on tailocin sensitivity (<u>Fig. 5</u>).

**Figure 5.**
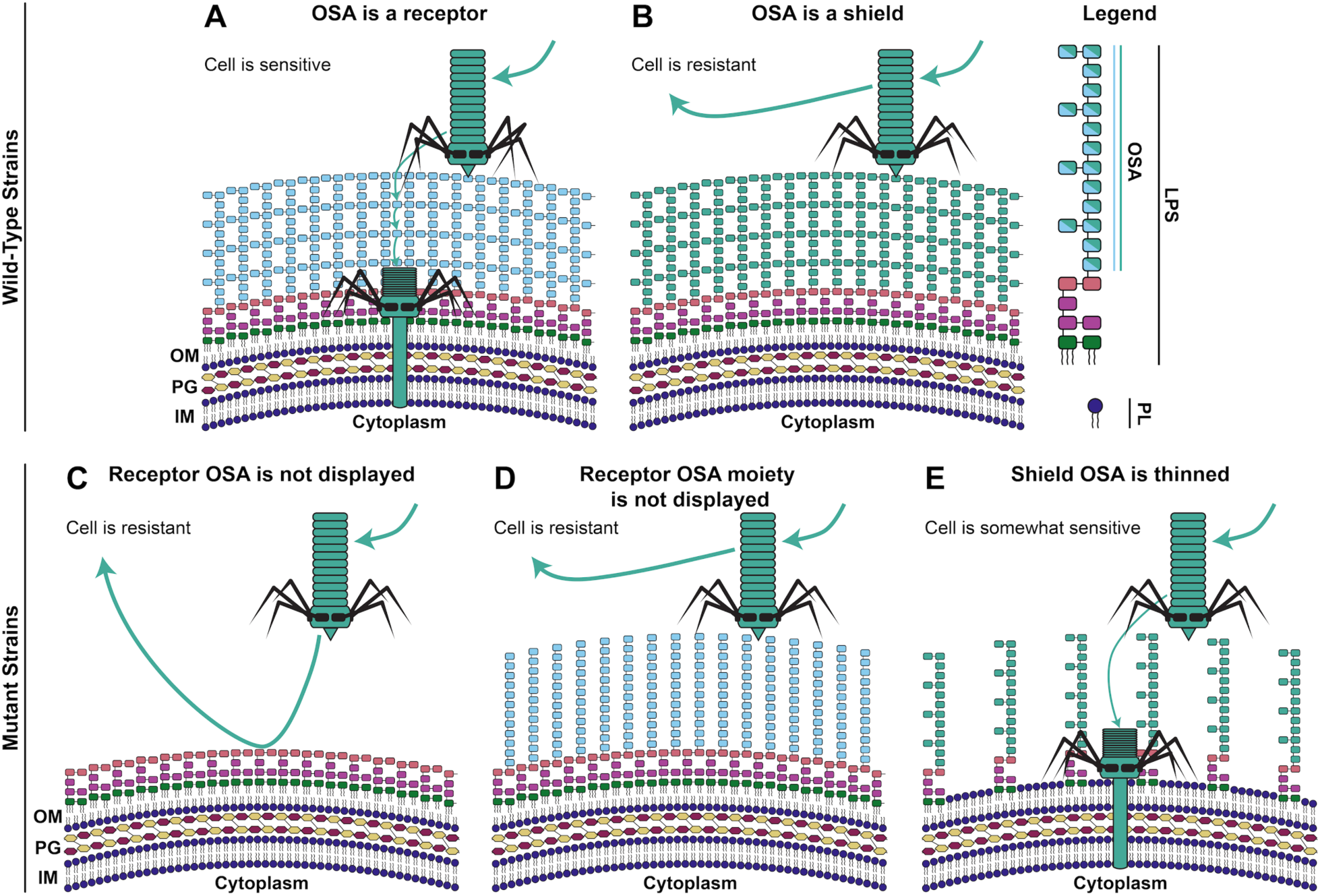
Proposed effect of O-specific antigens on tailocin lethality. We summarize our high-throughput reverse genetics data to propose 5 different OSA display scenarios that define whether a strain is sensitive or resistant to a tailocin. **(A)** Strain displays an LPS molecule whose OSA can be bound by the tailocin (is a receptor). The strain is naturally sensitive. **(B)** Strain displays an LPS molecule whose OSA interferes with tailocin binding (is a non-receptor/shield). The strain is naturally resistant. **(C)** Mutation in OSA biosynthesis/display leads to inability of the mutant to display receptor OSA. The mutant has become resistant. **(D)** Mutation in OSA biosynthesis/display leads to inability of the mutant to display the specific receptor OSA moiety. The mutant has become resistant. **(E)** Mutation in retrograde phospholipid transport or anterograde LPS transport leads to a thinning of the mutant’s shield LPS. The mutant has become more sensitive to tailocins. OM, outer membrane; PG, peptidoglycan; IM, inner membrane; OSA, O-specific antigen; LPS, lipopolysaccharide; PL, phospholipid.

Our data supports a model put forward by Köhler et al. (2010)^13^ that LPS–or, specifically, the OSA–acts as either a receptor for, or shield against, tailocins. Receptor OSA molecules can be bound by the tailocins, recruiting them toward the cell envelope, increasing the likelihood of lethal penetration, resulting in sensitivity. Shield OSA molecules act as a physical barrier between the tailocin and the cell envelope, resulting in resistance. Mutation resulting in removal of the entire receptor OSA, or just the specific receptor moiety of the OSA, can make a sensitive strain resistant. Alternatively, if the strain already possesses a shield OSA against the tailocin, that shield can be weakened by thinning the density of OSA at the cell envelope, in turn increasing sensitivity to tailocins. In other words, the composition and density of the LPS can help protect bacteria from tailocins, just as it protects them from small molecule antimicrobials and phage infections^54^.

Assumptions made under this model (<u>Fig. 5</u>) include: 1) uncapped LPS alone can act as a sufficient shield against our tailocins; and 2) tailocins can kill any strain, so long as they can get close enough to the cell envelope to penetrate completely. The first assumption is based on evidence that many phages (including HK620, P22, Sf6, 9NA and Det7) are unable to infect strains displaying an uncapped (OSA lacking) LPS^55^. The second assumption is made from the observation that wild-type Pse06, which is ‘resistant’ to its own tailocins, still displays an impaired OD_600_ growth curve in the presence of those tailocins (<u>Fig. 4C</u>). This observation could be explained by low frequency killing of the producing strain by its own tailocins should they spontaneously pass through the OSA shield and contract close enough to the cell envelope. Thus, we perhaps need to be careful how we use the terms ‘sensitive’ and ‘resistant’, which are assigned upon the presence or absence respectively of a zone of inhibition during a spot assay. Instead, intermediary phenotypes may be common, and may depend on genetic and temporal variation in LPS display and packing. Acquisition of more nuanced data could be achieved through widespread use of growth curves in assessing tailocin sensitivity in future.

Since our RB-TnSeq fitness assays yield a small set of very high scoring genes, identification of the most important factors involved in tailocin sensitivity is straightforward. However, strong positive selection for mutations in these genes shrouds our ability to detect factors with weaker impact on tailocin sensitivity. Mutants with lower relative fitness are rapidly outcompeted from the population and become undetectable in our data. Furthermore, we recognize that RB-TnSeq limits us to the study of non-essential genes. Many genes that modify the cell envelope (e.g. LPS and PL biosynthetic and transport genes) are essential, as noted in <u>Fig. 2C</u> and <u>Fig. 3B</u>. We were unable to evaluate the impact of these and other essential genes on tailocin sensitivity in this study. Future studies could improve on this using complementary mutational approaches^56,57^ that have recently been applied to investigate the genetic determinants of phage sensitivity^35^.

Overall, our fitness data suggests that tailocin sensitivity factors are limited to those affecting LPS composition and display, and can be found at well-defined genomic loci known to influence these phenotypes. The fitness data specifically emphasizes the importance of the OSA. However, this assertion is somewhat disputed by our high-throughput tailocin sensitivity phenotyping data, in which a subset of strains can vary in tailocin sensitivity despite sharing very similar OSA biosynthetic clusters. In our SM strains, differences in tailocin sensitivity cannot be explained by nucleotide variation in the OSA cluster (<u>Fig. S9</u>), *mla* or *lpt* genes, or by GWAS analysis. Thus, the responsible factors may be hard to discern at the sequence level, or may have resulted from sequence changes after we sequenced the strains. Gammaproteobacteria are known to escape phage predation through reversible and heritable (i.e. phase variable) changes in the expression of LPS modifying proteins^58–63^. It is possible that similar mechanisms are at work in our SM strains. In summary, the genotypes of tailocin sensitivity are more complicated than just the presence/absence of LPS-modifying genes.

This study represents the most comprehensive examination of tailocin sensitivity determinants to date. We assign new experimental tailocin sensitivity phenotypes to orthologs of both well-studied and hypothetical genes, highlighting the efficacy of our methods for studying non-model bacteria. We also perform the first investigation of resistance to tailocin-mediated self-intoxication and indicate the importance of LPS density in protecting a strain from its own tailocins. Our findings give weight to a model that LPS acts as either a receptor for, or a shield against, tailocins^13^, and provide a framework for studying bacterial evolution in the context of tailocin-mediated interference competition.

## Methods

### Bacterial strains, oligos, plasmids, and growth conditions

Strains used in this study are described in <u>Table S1</u>, oligos in <u>Table S2</u> and plasmids in <u>Table S3</u>. All cultures were aerobic. LB-Lennox (LB-L) media^68^ (Sigma-Aldrich) was used to cultivate all bacteria. *E. coli* and *P. aeruginosa* were incubated at 37ºC. *P. fluorescens, P. syringae* and groundwater *Pseudomonas* isolates were incubated at 30ºC. Liquid cultures were shaken at 200rpm, except for 96-well plate cultures which were shaken at 750rpm. When appropriate, antibiotics were supplemented to LB-L cultures at the following concentrations: carbenicillin at 100µg/mL (LB-L-Cb), kanamycin at 50µg/mL (LB-L-Km).

### Strain isolation

All environmental strains described in this study were isolated from groundwater wells at the Oak Ridge Field Research Center in Oak Ridge, TN. We used 1mL aliquots of groundwater to inoculate various types of liquid media for growth. Positive growth was identified by an increase in culture turbidity. Positive cultures were sequentially transferred in the same media twice, before streaking onto agar plates for single colonies. Individual colonies were picked and restreaked again to check purity. For storage, axenic colonies were regrown in liquid media to mid-log phase, amended with sterile glycerol to a final concentration of 30%, flash frozen with liquid nitrogen, and stored at −80ºC. The groundwater well, media and growth conditions used to isolate each strain can be found in <u>Table S1</u>^23,69–72^. While a variety of aerobic and anaerobic enrichment strategies were used for isolation, all strains are capable of growth in LB-L aerobically.

### Genome sequencing

For genomic DNA extraction, isolates were revived from glycerol stocks into 500µL liquid LB-L media in wells of 2.0mL 96-well DeepWell™ blocks (ThermoFisher™ Nunc™). The blocks were incubated with shaking until the cultures reached stationary phase. Cells were pelleted by centrifugation (3220g, 5mins) and the supernatant was decanted. Genomic DNA was manually purified using the QIAamp 96 DNA QIAcube HT kit (QIAGEN) and a vacuum manifold. All samples were eluted in AE buffer (10mM Tris-HCl, 0.5mM EDTA, pH 9.0). The samples were then randomly plated into a 384-well plate for automated Illumina library preparation. Each genomic DNA sample was normalized to 0.2ng/µL in 10mM Tris (pH 8.0). Then, libraries were prepared using the Nextera XT kit (Illumina) at 1/12th reaction size with a TTP LabTech mosquito^®^ HV liquid handling robot. Final libraries were cleaned with SPRIselect beads (Beckman Coulter) and pooled before sequencing on an Illumina NextSeq 500 producing 2×150bp paired-end reads. Separately, genomic DNA from the 12 Pse## codenamed strains (<u>Table S1</u>) was library prepped and sequenced on a PacBio Sequel at the Vincent J. Coates Genomics Sequencing Laboratory at UC Berkeley. Standard protocols were used for PacBio library prep and the PacBio reads were combined with Illumina reads for assembly.

### Genome assembly

Reads were preprocessed using BBtools version 38.60 to remove Illumina adapters, perform quality filtering and trimming, and remove PhiX174 spike-ins. We are not aware of any published papers documenting these tools. However, it is a standard tool suite developed at the Department of Energy Joint Genome Institute (JGI) and is documented at https://jgi.doe.gov/data-and-tools/bbtools/. Processing was done in two passes. First bbduk.sh was run with parameters *ktrim=r k=23 mink=11 hdist=1 ref=adapters*.*fa tbo tpe 2*. This was to remove any remaining Illumina adapters given in adapters.fa (standard Illumina adapters). Then bbduk.sh was run again with parameters *bf1 k=27 hdist=1 qtrim=rl trimq=17 cardinality=t ref=phix174_Illumina*.*fa*. This was to perform quality filtering and trimming as well as remove Illumina PhiX174 spike-ins given in the file phix174_Illumina.fa. Assembly was performed using SPAdes version 3.13.0^73^ with parameters *-k 21,33,55,77,99,127*.

### Identification of tailocin clusters

Tailocin elements were identified in assemblies of the studied isolates and in selected complete *Pseudomonas* spp. genomes using the PHASTER web-server^30^. Groups of orthologous genes (orthogroups) were identified using OrthoFinder 2.0^52^. Insertion sites for tailocin loci were identified by searching for orthologs of *mutS, cinA, trpE*, and *trpG* genes. A locus between *mutS* and *cinA* was considered tailocin-encoding if PHASTER identified it as an incomplete prophage. No tailocin-encoding genes were found between *trpE* and *trpG* genes in any of the studied isolates. Functional annotations for tailocin genes were determined by transfer from orthologous tailocin genes previously characterized in *P. chlororaphis*^*32*^, *P. putida* and *P. aeruginosa*^*26*^. For genes lacking characterized orthologs, functions were assigned in accordance with PHASTER annotations. Tailocin clusters were visualized using the R package ‘genoPlotR’^74^.

### Induction and purification of tailocins

5mL *P. fluorescens* LB-L cultures were cultivated overnight. These were then back-diluted 1:100 in 100mL LB-L and incubated with shaking in baffled 500mL Erlenmeyer flasks until OD_600_ reached ∼0.5. At this point, mitomycin C was applied to a final concentration of 0.5µg/mL to induce tailocin production, then incubation with shaking was resumed for an additional ∼18hrs. 10µL chloroform was added to lyse remaining intact cells. Cultures were centrifuged (3,220g, 1hr, 4ºC) to pellet cell debris. Supernatant was then sterilized by filtration (0.22µm filter). Ammonium sulfate was added to 30% saturation (16.4g/100mL), and dissolved by stirring with a magnetic stir bar in the cold room for 30mins. Precipitate was collected by centrifugation (16,000g, 37.5mins, 4ºC). The supernatant was decanted, and the precipitate was resuspended in 4mL SM buffer (100mM NaCl, 8mM MgSO_4_•7H_2_O, 50mM Tris-Cl, 0.01% gelatin) and stored at 4ºC. Uninduced control samples were prepared in the same way as experimental samples except that mitomycin C addition was omitted.

### Low-throughput spot test phenotypic assay

25×150mm glass culture tubes of 3mL liquid LB-L media were each inoculated with a different target strain from a frozen glycerol stock. The tubes were incubated with shaking until the cultures reached stationary phase. In the meantime, 100×15mm Petri dishes (BD Biosciences) were prepared with ∼25mL standard LB-L (1.5%) agar and cooled. 200µL of each stationary phase culture was mixed with 5mL molten LB-L soft (0.5%) agar, then immediately decanted over a pre-poured 1.5% LB-L agar plate with care taken to cover the entire surface. The plates were allowed to cool for 30mins at room temperature. 2µL stock or serially diluted tailocin sample was pipetted onto the solidified soft agar, then allowed to dry for 10mins. The plates were then incubated, and the interaction was deemed sensitive if a zone of inhibition was observed within 40hrs. When testing serially diluted tailocin samples, no plaques were observed, only zones of inhibition. This indicates that the killing agent is nonreplicative, and thus not reminiscent of a phage. When testing uninduced tailocin control samples (prepared without mitomycin C application) no lethality was observed (data not shown). Spot tests were performed in biological triplicates.

### Higher-throughput spot test phenotypic assay

Wells of 2.0mL 96-well DeepWell™ blocks (ThermoFisher™ Nunc™) were filled with 1mL liquid LB-L media, then each well was inoculated with a different target strain from a frozen glycerol stock. The blocks were incubated with shaking until the cultures reached stationary phase. In the meantime, 48-well Bio-One CELLSTAR^®^ plates (Greiner) were prepared with 800µL standard LB-L (1.5%) agar in each well, and cooled. 6µL of each stationary phase culture was mixed with 160µL molten LB-L soft (0.5%) agar, then immediately pipetted into a well of a pre-prepared 1.5% LB-L agar 48-well plate with care taken to cover the entire surface. The plates were allowed to cool for 30mins at room temperature. 2µL stock or serially diluted tailocin sample was pipetted onto the soft agar, then allowed to dry for 10mins. The plates were then incubated, and the interaction was deemed sensitive if a zone of inhibition was observed within 40hrs.

### Transmission electron microscopy (TEM)

TEM was performed at Donner Laboratory at Lawrence Berkeley National Laboratory. 4µL of tailocin samples, purified as above, were placed on glow-discharged carbon-coated grids (Formvar-carbon, 200 mesh copper, Electron Microscopy Sciences) for 5mins, then partially blotted to ∼1µL. Grids were then placed sample side down on a drop of ddH_2_O for 5mins, then transferred to a second drop of ddH_2_O for another 5mins. Residual water was partially blotted and 3µL of 2% aqueous uranyl acetate was applied. After 2mins, the sample was blotted to dryness. Grids were visualized on a JEOL JEM-1200x electron microscope operating at an accelerating voltage of 80kV. Images were recorded at 30,000X and 60,000X magnification with a charge-coupled-device (CCD) camera (UltraScan^®^, Gatan). Image analysis was done using ImageJ.

### Protein mass spectrometry

Tailocins purified as above were subjected to tryptic digestion and carboxyamidomethylation of cysteines performed in 40% (v/v) methanol, 5mM TCEP, 100mM ammonium bicarbonate, at pH 8.5. Mass spectrometry was then performed by the UC Berkeley Vincent J. Coates Proteomics/Mass Spectrometry Laboratory. MudPIT methods were used in order to achieve good sequence coverage of target proteins in a complex mixture^75^. A nano-LC column was packed in a 100µm inner diameter glass capillary with an emitter tip. The column consisted of 10cm of Polaris c18 5μm packing material (Varian), followed by 4cm of Partisphere 5 SCX (Whatman). The column was loaded by use of a pressure bomb and washed extensively with buffer A (5% acetonitrile and 0.02% heptafluorobutyric acid). The column was then directly coupled to an electrospray ionization source mounted on a Thermo-Fisher LTQ XL linear ion trap mass spectrometer. An Agilent 1200 HPLC equipped with a split line so as to deliver a flow rate of 300nL/min was used for chromatography. Peptides were eluted using a 4-step MudPIT procedure^75^.

### Analysis of mass spectrometry data

Protein identification was done using the Integrated Proteomics Pipeline (IP2, Integrated Proteomics Applications, Inc. San Diego, CA) using ProLuCID/Sequest, DTASelect2 and Census^76–79^. Tandem mass spectra were extracted into ms1 and ms2 files from raw files using RawExtractor^80^. Data was searched against a database of protein sequences specific for each sample supplemented with sequences of common contaminants and concatenated to a decoy database in which the sequence for each entry in the original database was reversed^81^. LTQ data was searched with 3000.0mmu precursor tolerance and the fragment ions were restricted to a 600.0ppm tolerance. All searches were parallelized and searched on the Vincent J. Coates proteomics cluster. Search space included all fully tryptic peptide candidates with no missed cleavage restrictions. Carbamidomethylation (+57.02146) of cysteine was considered a static modification. We required 1 peptide per protein and both tryptic termini for each protein identification. The ProLuCID search results were assembled and filtered using the DTASelect program^77,78^ with a peptide false discovery rate (FDR) of 0.001 for single peptides and a peptide FDR of 0.005 for additional peptides for the same protein. The molar percentage of protein content for each predicted tailocin protein was calculated from exponentially modified protein abundance index (emPAI)^82^ as 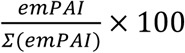 and is listed in <u>Table S7</u>. Overall, the following proportions of each tailocin sample are tailocin proteins: Pse04 Tailocins (15.6%); Pse05 Tailocins (9.4%); Pse06 Tailocins (7.7%), Pse11 Tailocins (14.6%).

### Competitive mutant fitness assays

We performed all our assays in 48-well microplates using the assay design, culture and data collection settings described previously^22^. In our assays, our partially purified tailocins were used as stressors at final concentrations 0.5x or 0.05x of the stock preparation. SM buffer was supplied as a tailocin-free control. We supplied LB-L (for wild-type strains) or LB-L-Km (for mutants) as the growth medium in each case. The microplates were incubated in a Tecan Infinite F200 plate reader with orbital shaking and OD_600_ readings every 15mins. Mutant library cultures were harvested for subsequent BarSeq analysis at mid-log phase, as determined from OD_600_ traces. Cell pellets were obtained by centrifugation (8,000g, 3mins) and stored at −20ºC awaiting genomic DNA extraction. Each condition was assayed in duplicate.

### BarSeq

Genomic DNA extraction and barcode PCR were performed as described previously^22,23^. All genomic DNA extractions were done using the DNeasy Blood and Tissue kit (Qiagen) and quantified using a NanoDrop 1000 device (Thermo Fisher). Thermocyclers were set to the 98ºC BarSeq PCR protocol (“BarSeq98”)^22^. A primer set that permits improved cluster discrimination on the Illumina HiSeq 4000 was used (<u>Table S2</u>)^23^. Barcode sequence data was obtained by multiplexing samples on a lane of a HiSeq 4000 (50-cycle single read) run at the UC Berkeley Vincent J. Coates Genomics Sequencing Laboratory. Fitness data was calculated and analyzed from BarSeq reads as previously described^22^.

### Analysis of BarSeq data

BarSeq data was analyzed as described previously^22^. Then, fitness hits were filtered further as in our phage resistance studies^35,36^ to reduce false positives caused by a lack of coverage of most mutants following stringent tailocin selection. To be evaluated as a positive fitness hit, gene disruptions had to exhibit a fitness score ≥7.0 (<u>Tables S10</u>, <u>S13</u> and <u>S16</u>), t-like statistic ≥5.0 (<u>Tables S11</u>, <u>S14</u> and <u>S17</u>) and show an increase in barcode read count from before treatment (Time0) to after treatment (<u>Tables S9</u>, <u>S12</u> and <u>S15</u>). To be evaluated as a negative fitness hit, gene disruptions had to exhibit a fitness score ≤-2.0 (<u>Table S19</u>), t-like statistic ≤-5.0 (<u>Table S20</u>), and show a decrease in barcode read count from before treatment (Time0) to after treatment (<u>Table S18</u>). Heat maps of fitness scores were generated using Morpheus (https://software.broadinstitute.org/morpheus).

### Validation of tailocin resistance phenotypes

Individual transposon-insertion mutants for validating tailocin resistance phenotypes were isolated by enriching them from their respective pooled RB-TnSeq libraries through the application of tailocins that the wild-type strain is sensitive to. RB-TnSeq libraries, at a starting OD_600_ of 0.02, were cultured with tailocins at 0.5x stock concentration, in the wells of a 96-well microplate (200µL per well). The microplates were incubated in a Tecan Infinite F200 plate reader with orbital shaking and OD_600_ readings every 15mins. Cultures were halted at early-log phase and spread on LB-L-Km agar for single colonies. Single colonies were subcultured in LB-L-Km, then subjected to colony arbitrary PCR to determine barcode and transposon insertion site. A description of the colony arbitrary PCR is as follows. 100µL saturated culture was pelleted, resuspended in 2.5% v/v IGEPAL^®^ CA-630 (Sigma-Aldrich) and boiled at 98ºC for 20mins. Cell debris was pelleted by centrifugation. 1µL supernatant was then used as template for the first round of arbitrary PCR: a 20µL total volume Q5^®^ Hot Start polymerase reaction (New England Biolabs) with 2.5pmol oALA051, 2.5pmol oALA052, and 5pmol oALA054 primers (see <u>Table S2</u> for primer sequences). First round arbitrary PCR thermal cycling conditions were: 98ºC for 3mins; 5 cycles of 98ºC for 30s, 42ºC (−1ºC/cycle) for 30s, 72ºC for 3mins; 25 cycles of 98ºC for 30s, 55ºC for 30s, 72ºC for 3mins; 72ºC for 5mins. 2µL first round PCR product was used as template for the second round of arbitrary PCR: 20µL total volume Q5^®^ Hot Start polymerase reaction with 2.5pmol oALA053, and 2.5pmol oALA055. Second round arbitrary PCR thermal cycling conditions were: 98ºC for 3mins; 30 cycles of 98ºC for 30s, 55ºC for 30s, 72ºC for 3mins; 72ºC for 5mins. 10µL second round arbitrary PCR product was cleaned using AMPure beads (Beckman Coulter) and Sanger sequenced using oALA055. 14 tailocin-resistant single strains were identified and isolated in this way (see <u>Tables S1 and S23</u>). Tailocin resistance was verified by spot test phenotypic assay (see above).

### Validation of tailocin production phenotypes

Tailocin production and tailocin sensitivity phenotypes were validated through marked deletions of baseplate genes and genes with significant negative fitness scores respectively. To do this, we assembled derivatives of the plasmid pMO7704 to generate replacements of our genes of interest with a kanamycin resistance marker via double crossover. Plasmid assembly and strain construction details can be found in <u>Tables S3 and S1</u> respectively. Plasmids propagated in the *E. coli* 10-beta cloning strain were purified using the standard QIAprep protocol (QIAGEN), then delivered to the *Pseudomonas* strain via previously published electroporation^83^ or conjugation^84^ protocols. Transformants were selected on LB-L-Km agar. Disruption of tailocin production was verified by spot test phenotypic assay (see above).

### Validation of tailocin sensitivity phenotypes

Increases in tailocin sensitivity were too subtle to be detected by spot assay (data not shown). Instead, sensitivity was verified by comparing growth in planktonic culture of mutants vs. wild-type in the presence of tailocins. Cultures were prepared at a starting OD_600_ of 0.02, with tailocins added at 0.5x stock concentration, in the wells of a 96-well microplate (200µL total volume per well). The microplates were incubated in a Tecan Infinite F200 plate reader with orbital shaking and OD_600_ readings every 15mins.

### Identification and comparison of lipopolysaccharide biosynthetic gene clusters

Gene clusters encoding LPS core oligosaccharide, O-specific antigen (OSA) and common polysaccharide antigen (CPA) biosynthetic enzymes were identified by analysis of orthologous groups of genes (orthogroups) as described above, using OrthoFinder 2.0^52^. Orthologs of *P. aeruginosa* PAO1 genes PA4997-PA5012 and PA5447-PA5459 were considered to represent LPS core oligosaccharide and CPA gene clusters, respectively, in accordance with existing data on gene functions^34^. As gene content of OSA biosynthetic loci is variable, highly conserved flanking genes were used to determine locus boundaries: *himD*/*ihfB* on the 5’ end and *wbpM* on the 3’ end^85^. All genes located between these two genes were considered to form OSA biosynthetic clusters. Gene clusters were visualized using the R package ‘genoPlotR’^74^. To compare gene contents of OSA clusters, Jaccard distances for all pairs of strains were computed as 1 minus the number of orthogroups present in OSA clusters of both strains divided by the number of orthogroups present in either of the OSA clusters. The resulting matrix was used to compute a hierarchical cluster using the “hclust” function in R (v3.0.2) (method=“single”) and visualized with the “dendextend” package in R and the Interactive Tree of Life (iTOL) online tool^86^.

### Phylogenetic analysis

To estimate phylogenetic relationships between genomes of our collection of *Pseudomonas* spp. strains, we identified a set of 120 bacterial marker genes with GTDB-Tk toolkit^24^. Only 88 marker genes were found in single copy in each of the 130 genomes studied. Protein sequences of those 88 markers were aligned by MAFFT v7.310^87^ with *--auto* option, and the resulting 88 alignments were concatenated into a single multiple sequence alignment. A phylogenetic tree was reconstructed from the multiple alignment using the maximum likelihood method implemented in the FastTree software v2.1.10^88^ and visualized using the Interactive Tree of Life (iTOL) online tool^86^.

## Author Contributions

V.K.M. conceived the project. A.P.A. and V.K.M. supervised the project. S.C. led the experimental work. S.C., A.L.A. and P.J.W. collected data. A.E.K. performed computational analysis. S.C., A.M.D., and V.K.M. analyzed the fitness data. M.N.P. and A.P.A. provided advice on data processing. R.C. isolated the bacterial strains. S.C., V.K.M., and A.P.A. wrote the paper.

## Acknowledgements

The authors acknowledge the work of ENIGMA Scientific Focus Area colleagues on isolating and sequencing the *Pseudomonas* spp. reported here, specifically: Alexander Aaring, Eric J. Alm, Mark Callaghan, Hans K. Carlson, Jennifer V. Kuehl, Lauren M. Lui, Ryan A. Melnyk, Torben Nielsen and Sarah J. Spencer. S.C. would like to acknowledge Benjamin A. Adler for discussions and critical feedback on the manuscript. This work was funded by ENIGMA, a Scientific Focus Area Program at Lawrence Berkeley National Laboratory, supported by the U.S. Department of Energy, Office of Science, Office of Biological and Environmental Research under contract DE-AC02-05CH11231. Proteomics data acquisition and analysis was performed at the Vincent J. Proteomics/Mass Spectrometry Laboratory at the University of California, Berkeley, supported in part by NIH S10 Instrumentation Grant S10RR025622. PacBio genome sequence and BarSeq data acquisition was performed at the Vincent J. Coates Genomics Sequencing Laboratory at the University of California, Berkeley, supported by NIH S19 Instrumentation Grants S10RR029668, S10RR027303, and OD018174. S.C. acknowledges support from: the NSF Graduate Research Fellowship Program under Grant No. DGE 1752814, the Croucher Foundation through a Croucher Scholarship for Doctoral Study, and the Drs Richard Charles and Esther Yewpick Lee Charitable Foundation through an R C Lee Centenary Scholarship. Any opinions, findings, and conclusions or recommendations expressed in this material are those of the authors and do not necessarily reflect the views of the National Science Foundation.

## Supplementary Information

### Supplementary Notes

- Supplementary Note 1. Review of the roles of LPS inner core biosynthetic enzymes on O-specific antigen display
- Supplementary Note 2. Putative functions of AO361_RS10865 and AO361_RS10900, genes involved in sensitivity to a subset of antagonistic tailocins

### Supplementary Tables (in separate .xlsx workbook)

- Table S1. Bacterial strains used in this study.
- Table S2. Oligonucleotide used in this study.
- Table S3. Plasmids used in this study.
- Table S4. Tailocin genes identified via our bioinformatics approach.
- Table S5. Tailocin gene orthogroups across *Pseudomonas* strains.
- Table S6. Killing activity of tailocin samples across 130 *Pseudomonas* strains.
- Table S7. Proteomics results for tailocin samples.
- Table S8. Description of RB-TnSeq mutant libraries used in this study.
- Table S9. Read count data for Pse05 genes implicated in tailocin sensitivity.
- Table S10. Additional fitness data for Pse05 genes implicated in tailocin sensitivity.
- Table S11. t-like statistics data for Pse05 genes implicated in tailocin sensitivity.
- Table S12. Read count data for Pse03 genes implicated in tailocin sensitivity.
- Table S13. Additional fitness data for Pse03 genes implicated in tailocin sensitivity.
- Table S14. t-like statistics data for Pse03 genes implicated in tailocin sensitivity.
- Table S15. Read count data for Pse13 genes implicated in tailocin sensitivity.
- Table S16. Additional fitness data for Pse13 genes implicated in tailocin sensitivity.
- Table S17. t-like statistics data for Pse13 genes implicated in tailocin sensitivity.
- Table S18. Read count data for Pse06 genes implicated in tailocin resistance.
- Table S19. Additional fitness data for Pse06 genes implicated in tailocin resistance.
- Table S20. t-like statistics data for Pse06 genes implicated in tailocin resistance.
- Table S21. LPS gene orthogroups across *Pseudomonas* strains.
- Table S22. O-specific antigen gene orthogroups across 13 *Pseudomonas* strains.
- Table S23. Descriptions of all enriched transposon-insertion mutants used for validations.
- Table S24. O-specific antigen gene orthogroups across 130 *Pseudomonas* strains.

**Figure S1.**
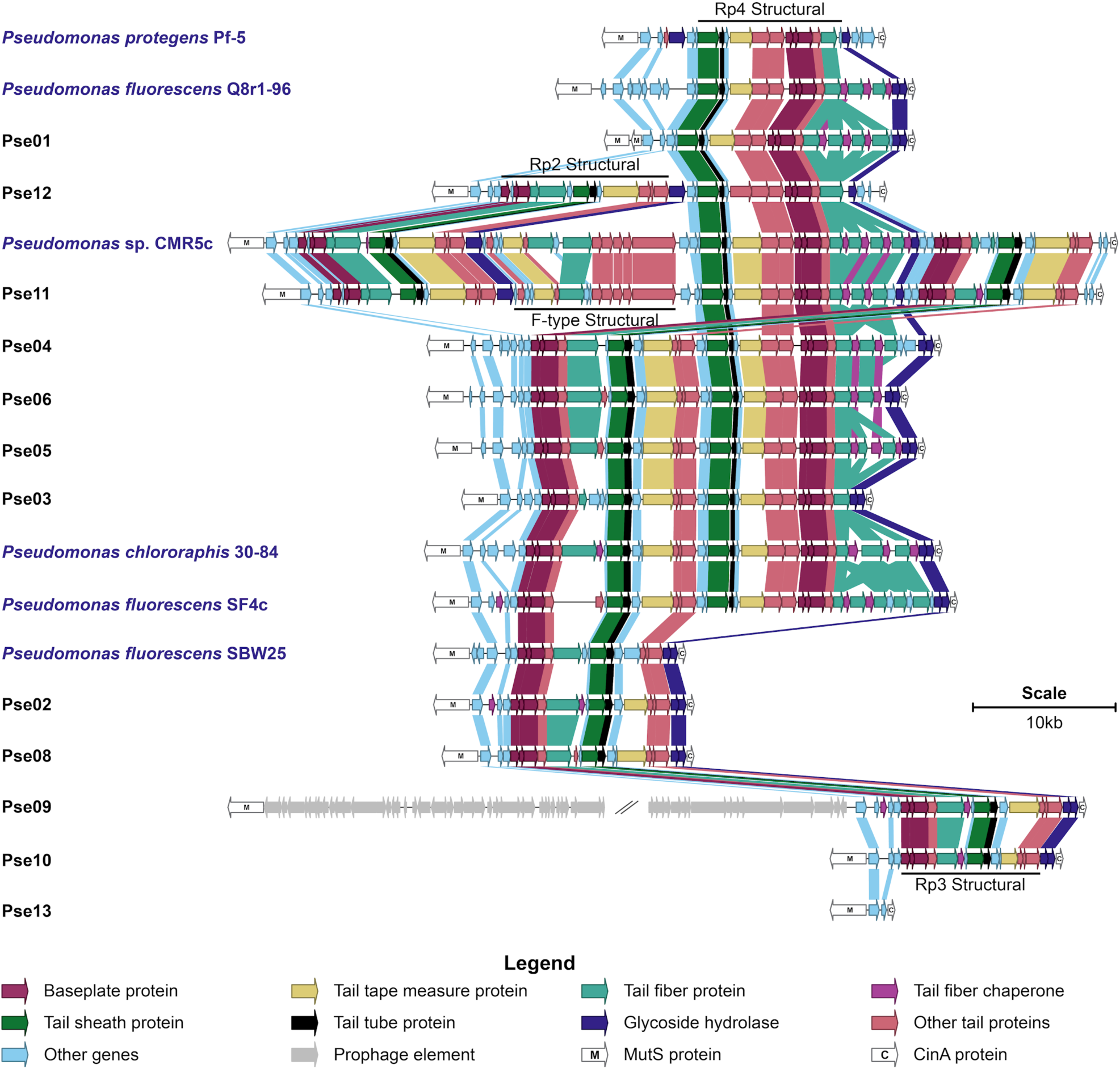
Extended tailocin gene clusters. Tailocin gene clusters of various sizes were found in 11 of our 12 selected *Pseudomonas* isolates, and solely at the *mutS*/*cinA* locus. Pse09 appears to encode a full prophage of unknown length at this locus in addition to an Rp3 tailocin cluster. Pse13 has no tailocin gene cluster. Tailocin gene clusters in previously studied soil and rhizosphere *Pseudomonas* strains (Pf-5, Q8r1-96, CMR5c, 30-84, SF4c, SBW25), labeled in blue text, are included for comparison. Genes are colored by the presence of key words in their PHASTER^30^ predicted annotations. Genes in the same orthogroup (<u>Methods</u>) are joined by a block of color. Additional data on these clusters can be found in <u>Table S4</u>. For a full list of orthogroups, see <u>Table S5</u>.

**Figure S2.**
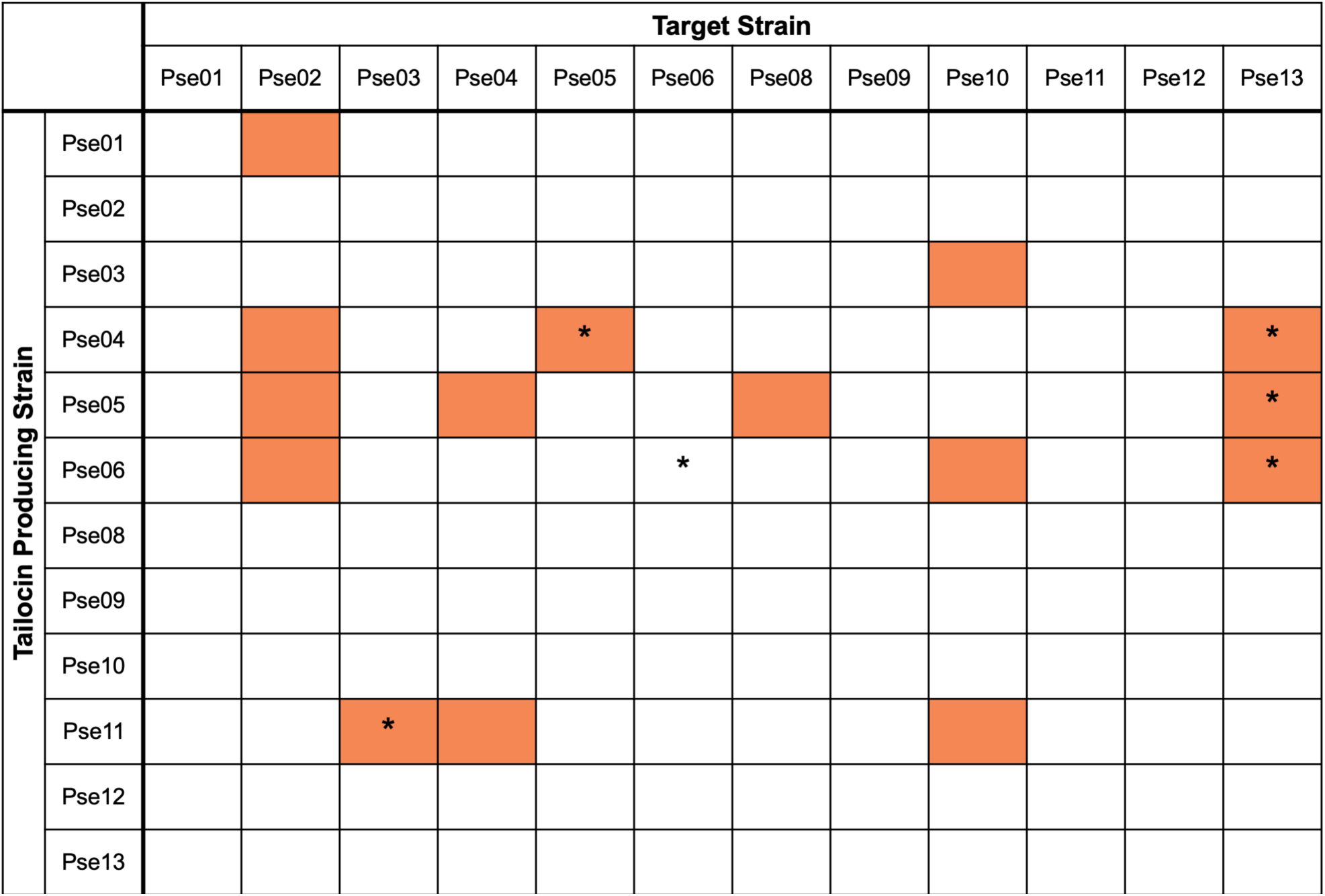
Initial tailocin susceptibility data. Partially purified tailocin samples from 12 *Pseudomonas* strains were used to challenge the same 12 strains by spot testing. Note: Pse13 does not encode tailocins, so we did not expect its ‘tailocin sample’ to be lethal to any target strain. Orange and white highlighted cells illustrate sensitive and resistant interactions respectively. No tailocin sample killed its producing strain. Starred (*) cells illustrate interactions that were investigated in greater depth due to the availability of RB-TnSeq^22^ mutant libraries in the target strains.

**Figure S3.**
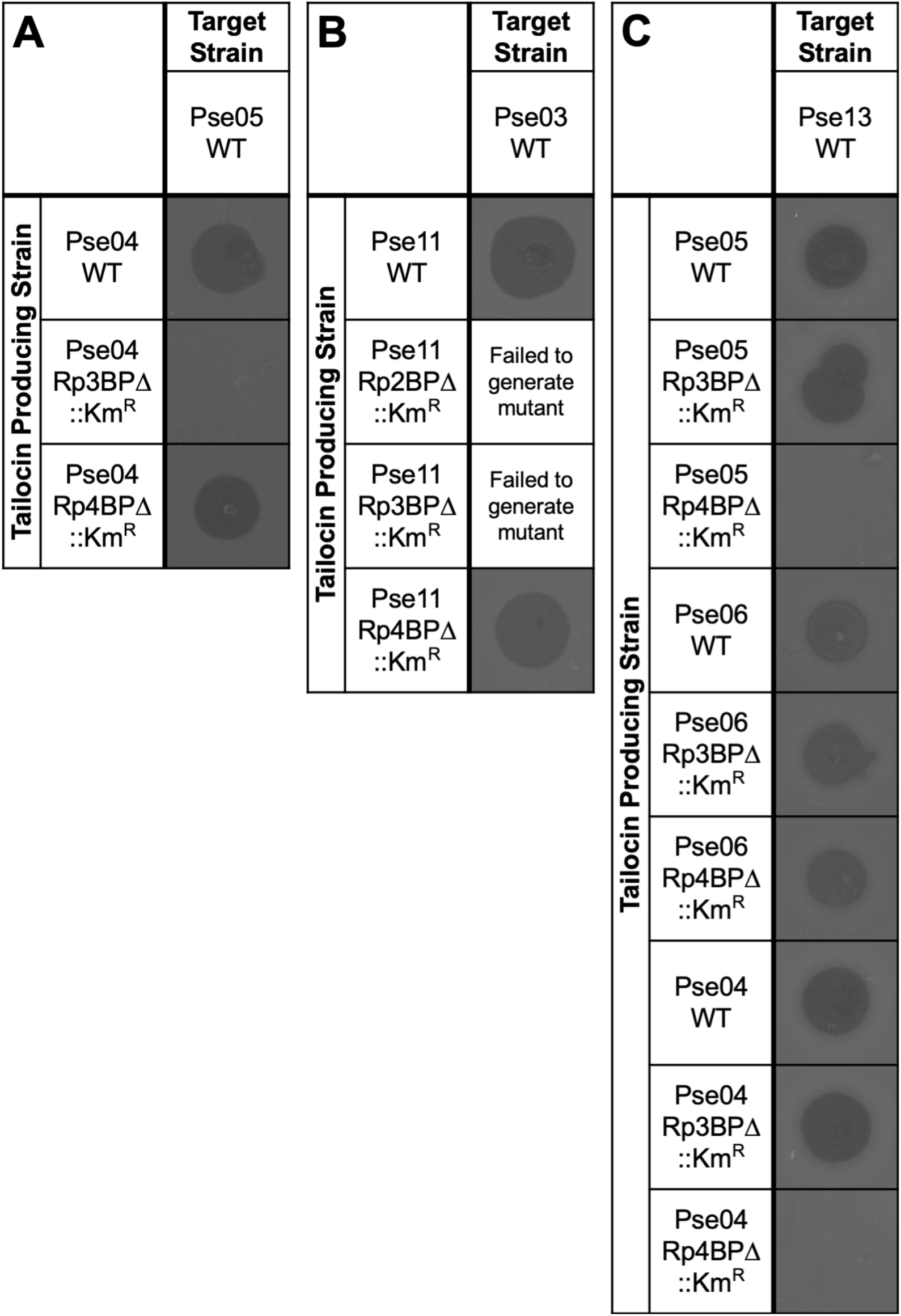
Tailocin production phenotype validations. Marked gene deletion vectors were used to replace baseplate protein genes with a kanamycin-resistance marker. Tailocins were induced and partially purified from these baseplate mutants and spotted on strains sensitive to the wild-type tailocin. Target strains are: (A) Pse05; (B) Pse03; (C) Pse13. BP, baseplate genes. This data suggests that the Pse04 Rp3 tailocin is responsible for killing Pse05, while its Rp4 tailocin in turn kills Pse13. Meanwhile, only the Rp4 tailocin of Pse05 appears to kill Pse13. Finally, both tailocin particles of Pse06 kill Pse13. We were unable to generate Rp2BP or Rp3BP mutants in Pse11.

**Figure S4.**
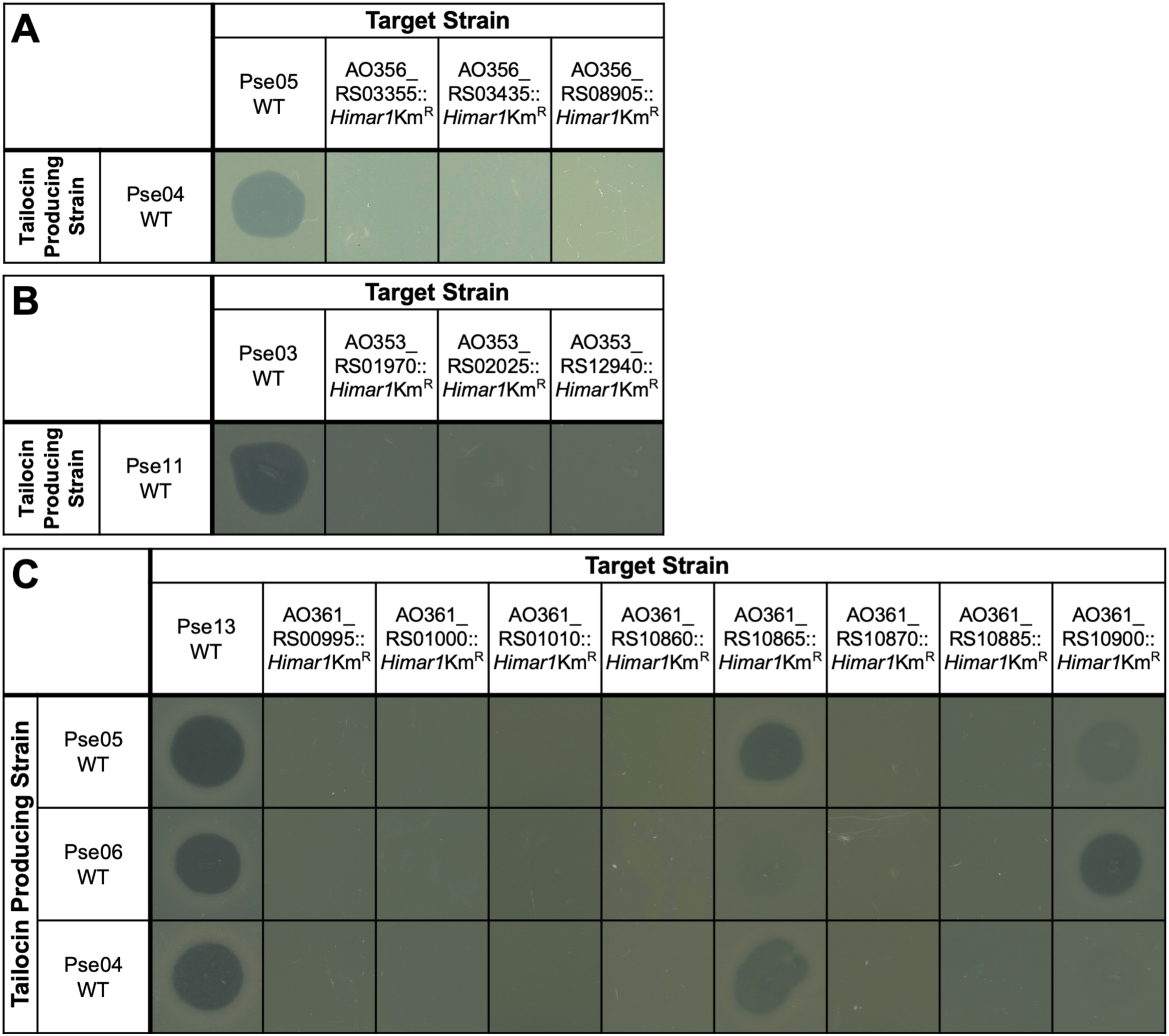
Tailocin resistance phenotype validations. Select RB-TnSeq mutants implied by our fitness data to be resistant to tailocins were isolated from their respective libraries and spotted with tailocin sample (see Methods). Target strains are: (A) Pse05 wild-type and mutants; (B) Pse03 wild-type and mutants; (C) Pse13 wild-type and mutants. An image of the presence or absence of a zone of clearance is shown. Some Pse13 mutants show differential sensitivity to tailocins from different producer strains. See <u>Table S23</u> for more information on these mutants.

**Figure S5.**
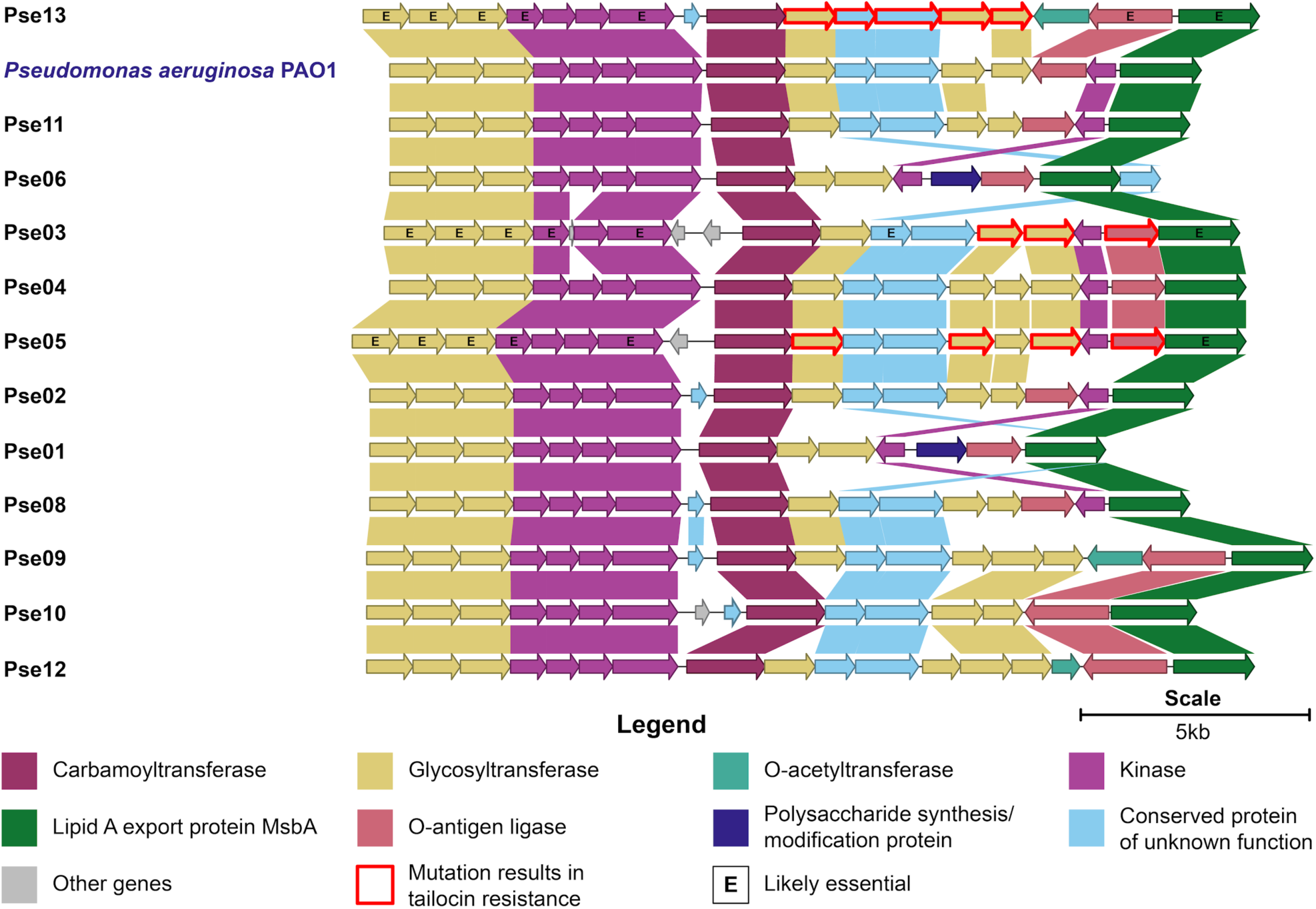
LPS core oligosaccharide biosynthetic gene clusters. Illustrated and compared are the genes comprising LPS core clusters encoded by our 12 selected *Pseudomonas* isolates (black) and *P. aeruginosa* PAO1 (blue label). These clusters show a high degree of homology. Genes in the same orthogroup (<u>Methods</u>) are joined by a block of color. For a full list of orthogroups, see <u>Table S21</u>.

**Figure S6.**
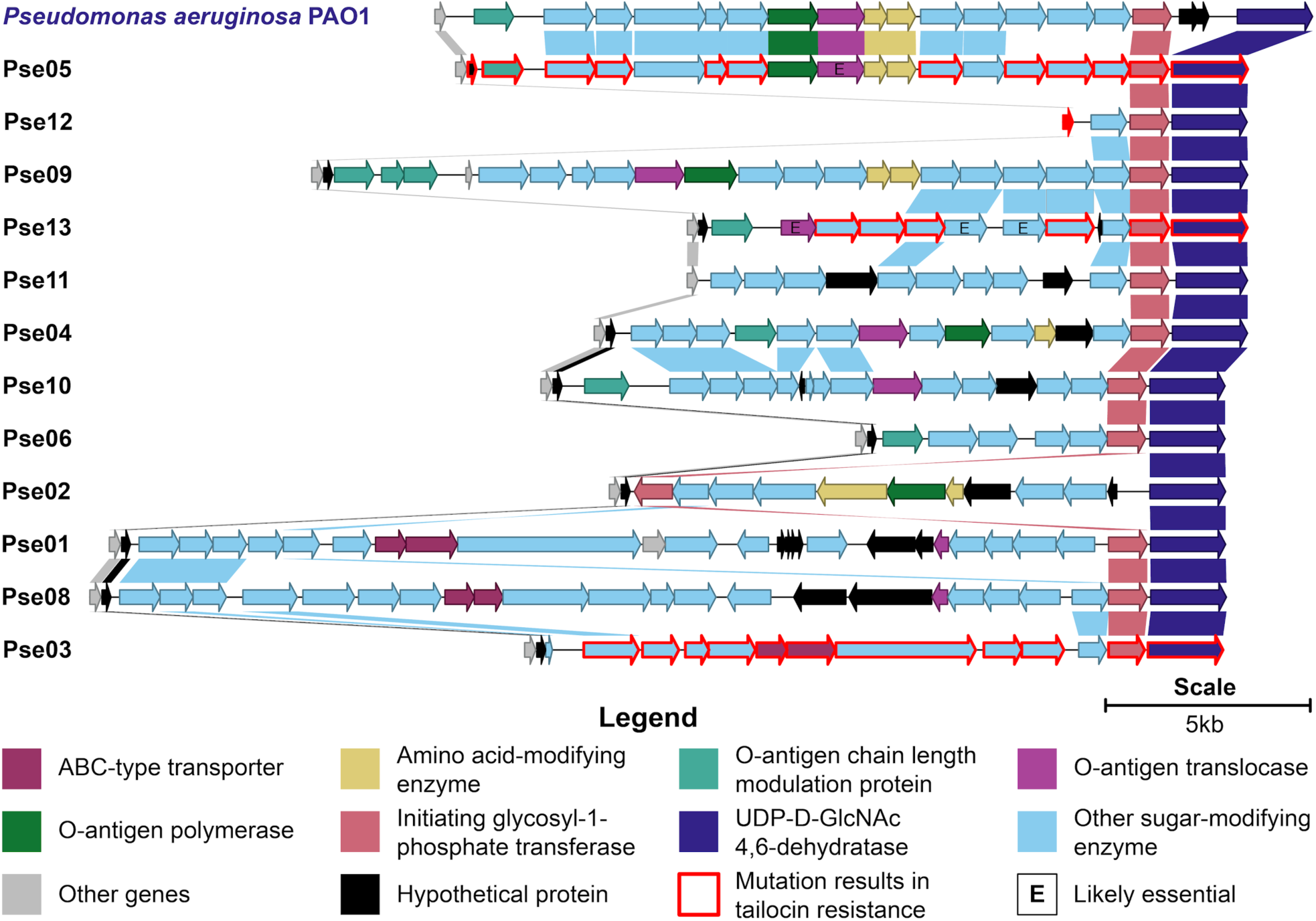
O-specific antigen biosynthetic gene clusters. Illustrated and compared are the genes comprising OSA biosynthetic clusters encoded by our 12 selected *Pseudomonas* isolates (black) and *P. aeruginosa* PAO1 (blue label). While these clusters display considerable heterogeneity, do note the high homology between the clusters of PAO1 and Pse05. Genes in the same orthogroup (<u>Methods)</u> are joined by a block of color. For a full list of orthogroups, see <u>Table S22</u>.

**Figure S7.**
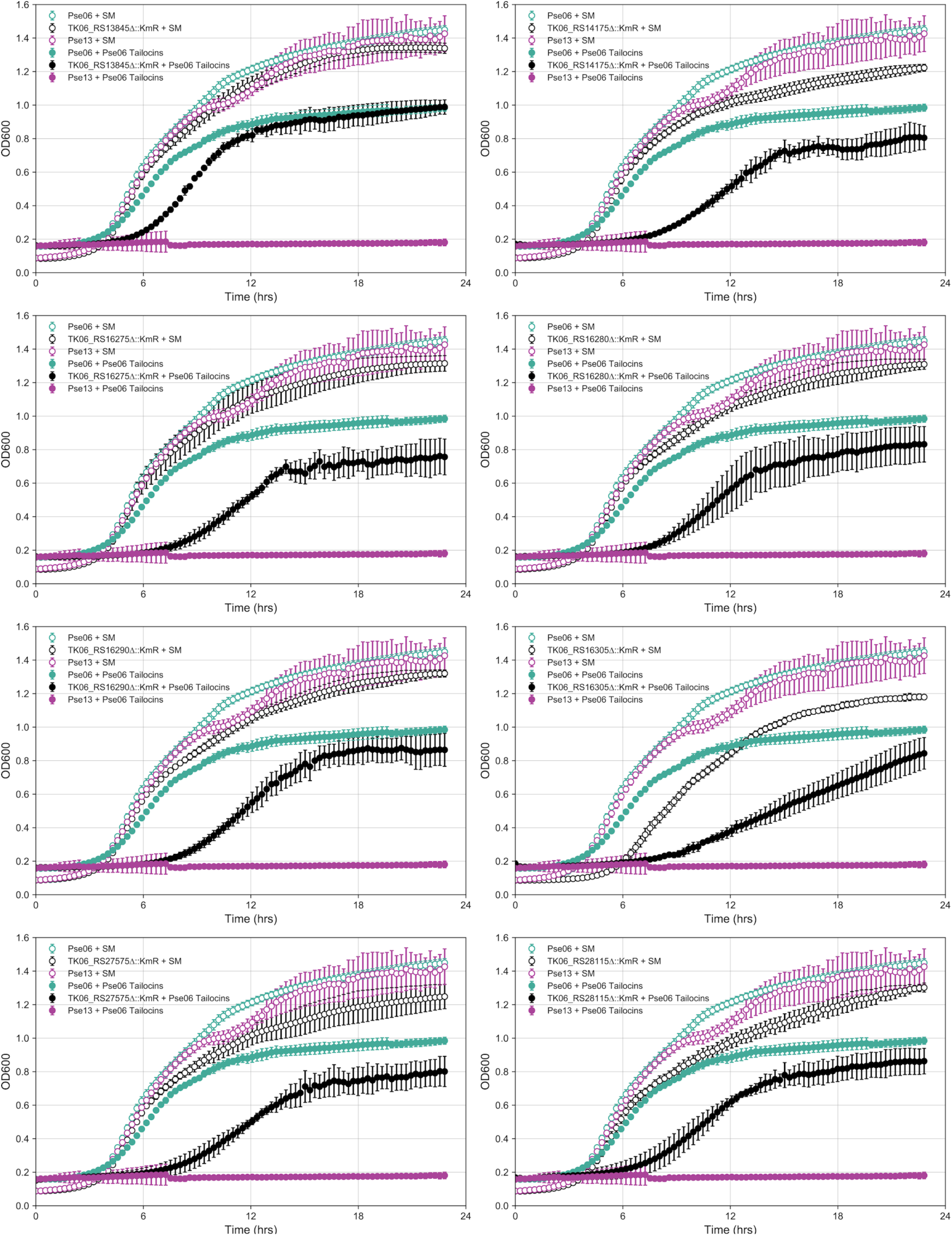
Tailocin sensitivity phenotype validations. Mutations in Pse06 that cause increased tailocin sensitivity were regenerated via replacement of the gene with a kanamycin-resistance marker. Growth curves of these mutants, wild-type Pse06 (naturally resistant) and wild-type Pse13 (naturally sensitive) were obtained in the presence and absence of Pse06 tailocins. Growth assays proceeded for 22.5hrs. For each mutant, application of Pse06 tailocins results in weaker growth compared to wild-type Pse06, but does not eliminate growth like with wild-type Pse13. These experiments were repeated (triplicate) and averages are plotted. Error bars: standard deviation.

**Figure S8.**
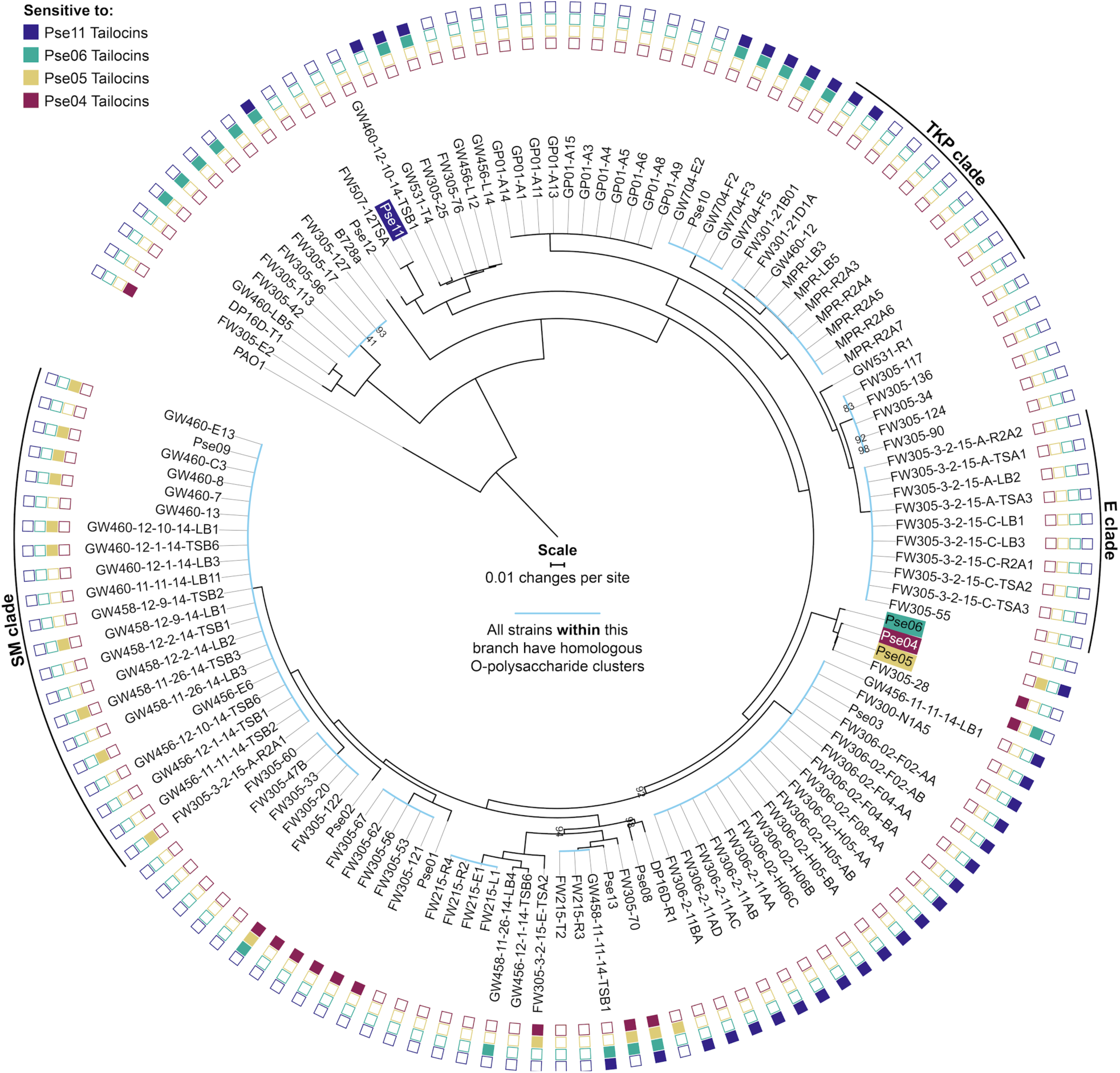
Phylogenetic tree of target strains overlaid with tailocin sensitivity data. GTDB-Tk (Genome Taxonomy Database Toolkit)^24^ was used to find 88 ubiquitous single-copy proteins found in each strain’s genome. Phylogenetic distances were then determined using a multilocus alignment of all 88 protein sequences (see <u>Methods</u>). The root of this tree is set to *P. aeruginosa* PAO1 in accordance with *Pseudomonas* phylogenomics^67^. All percentage bootstrap values <100% are labeled at branching nodes. Light blue colored branches indicate clades whose members all internally share a homologous O-specific antigen cluster, after we have annotated these clusters per <u>Methods</u>. Tailocin producing strain labels are highlighted in color. Shaded boxes at the outer edge of the tree indicate sensitivity of that strain to the correspondingly colored tailocin. To aid discussion of 3 clades of strains with interesting features, we named those clades after the most closely related species by 16S rRNA similarity: E clade (*P. extremorientalis*); TKP clade (*P*. sp. TKP); SM clade (*P. silesiensis*/*P*.*mandelii*). For the killing matrix in table format, see <u>Table S6</u>.

**Figure S9.**
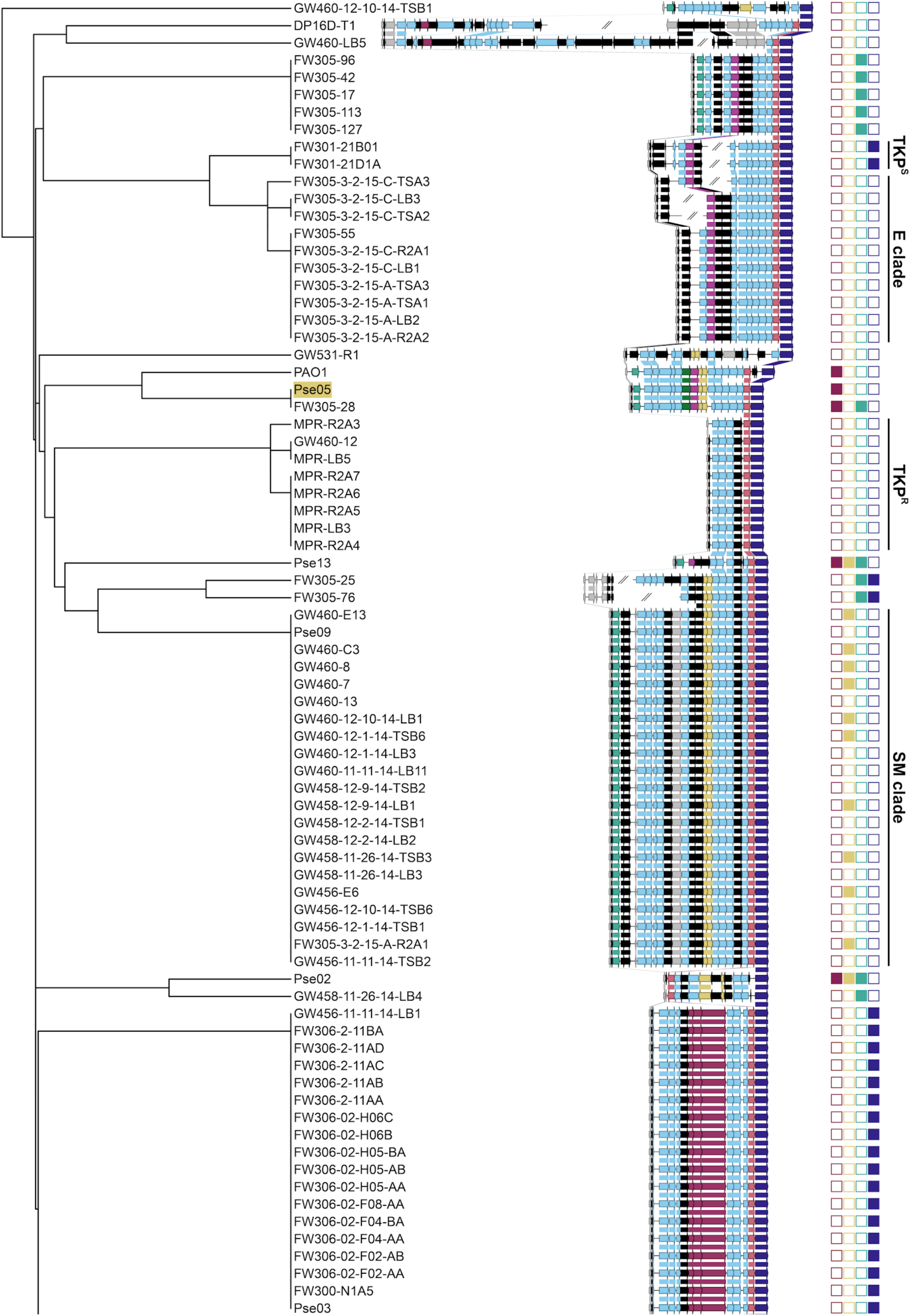
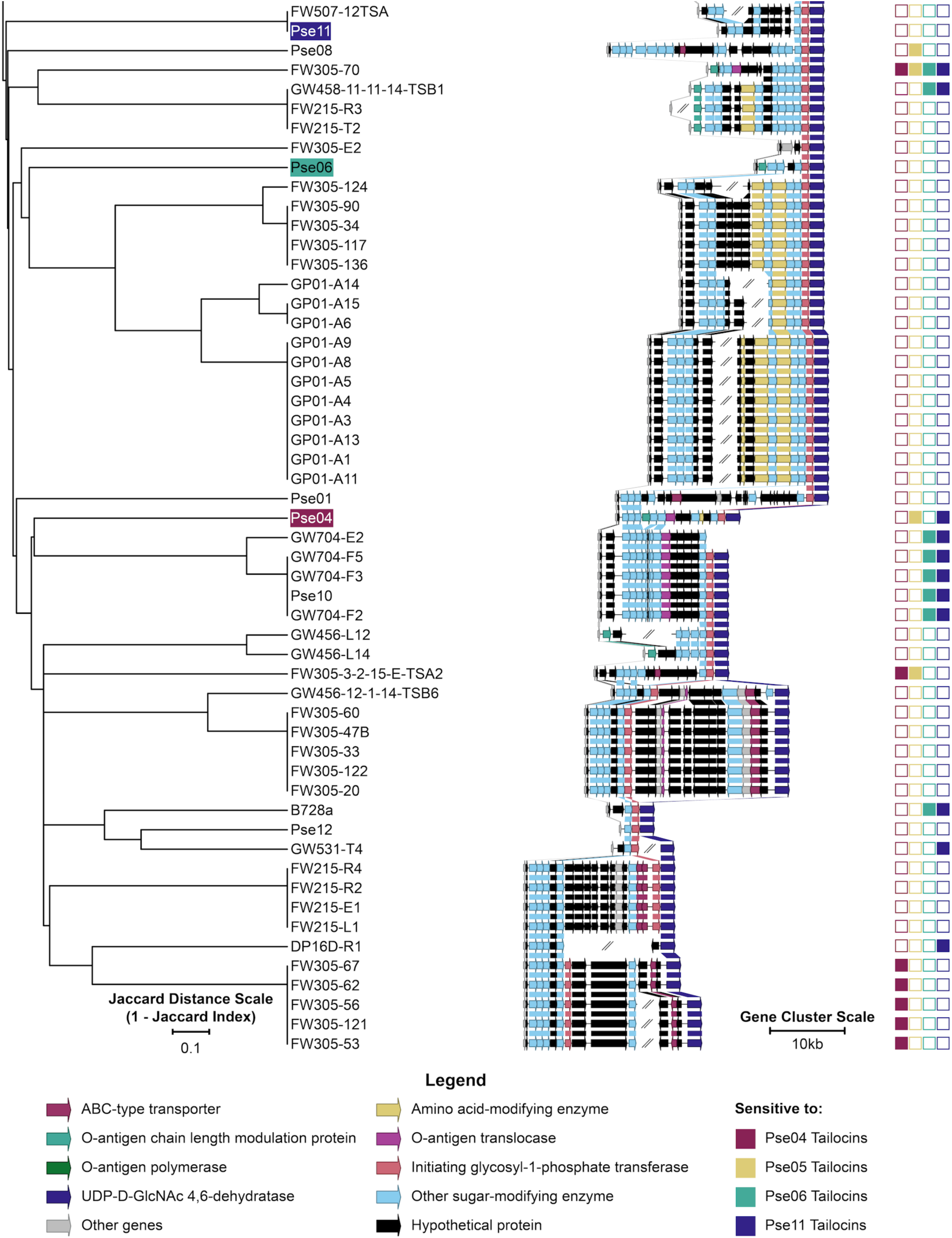
O-specific antigen biosynthetic gene clusters of target strains overlaid with tailocin sensitivity data. Illustrations of the OSA clusters of all 130 *Pseudomonas* strains clustered in the same way as shown in <u>Fig. 4</u>. OSA clusters are defined as the region flanked by the genes *ihfB* (integration host factor subunit beta, left, non-inclusive) and *wbpM* (UDP-D-GlcNAc 4,6-dehydratase, right, inclusive). Genes in the same orthogroup (<u>Methods</u>) are joined by a block of color. Tailocin producing strain labels are highlighted in color. Shaded boxes on the right indicate sensitivity of that strain to the correspondingly colored tailocin. To aid discussion of 3 clades of strains with interesting features, we named those clades after the most closely related species by 16S rRNA similarity: E clade (*P. extremorientalis*); TKP clade (*P*. sp. TKP); SM clade (*P. silesiensis*/*P. mandelii*). In this clustering, TKP strains are separated into two groups: TKP^S^ (Pse11 Tailocins sensitive) and TKP^R^ (resistant). For the killing matrix in table format, see <u>Table S6</u>. For a full list of orthogroups, see <u>Table S24</u>.

**Supplementary Note 1. Review of the roles of LPS inner core biosynthetic enzymes on O-specific antigen display**

Studies in *Escherichia coli* and *Salmonella enterica* show truncation of the LPS and elimination of OSA display following disruption of LPS inner core assembly genes (i.e. *waaA, waaC, waaF* and *waaG*)^54^. However, we are unable to evaluate their effect on tailocin sensitivity in this study, since all of these genes are essential in *P. aeruginosa* PAO1^34^ and likely essential (see <u>Methods</u>) in Pse05, Pse03 and Pse13. The one exception is *waaC* (AO353_RS12880) in Pse03, which confers neutral fitness to Pse11 tailocins when disrupted. This gene is disrupted just three times in the Pse03 RB-TnSeq library, with one transposon insertion at the 483rd base pair, and two insertions at the 531st base pair of this 1073bp gene. As the median number of insertions per gene in this library is eleven^23^, we postulate that the three non-lethal insertions in AO353_RS12880 may not have disrupted its essential LPS core assembly function, leaving O-specific antigen display unaffected. A viable transposon-insertion in *waaC* has been reported once before, in PAO1, in the 1038(1068)bp position, so this gene may tolerate select insertion sites^64^.

**Supplementary Note 2. Putative functions of AO361_RS10865 and AO361_RS10900, genes involved in sensitivity to a subset of antagonistic tailocins** AO361_RS10865 encodes a putative glycosyltransferase but shares no significant similarity to any characterized enzyme. AO361_RS10900 is an ortholog of *wbpM* in PAO1 (78% identity, 100% coverage) and may share with *wbpM* a function in modifying OSA monomers. In PAO1, *wbpM* encodes a bifunctional, inner-membrane-bound sugar C6 dehydratase/C4 reductase that catalyzes the conversion of UDP-D-GlcNAc to UDP-4-keto-D-QuiNAc^65^. This is an intermediary step in the biosynthesis of D-QuiNAc or D-FucNAc, monomers in the O-specific antigen (OSA) of *P. aeruginosa* serotypes O3, O5, O6 and O10. Deletion of *wbpM* in serotype O3, O5, O6 and O10 results in complete loss of the OSA. In contrast, deletion of *wbpM* in serotypes O15 and O17 results in slight modification to the OSA, as determined by silver-stained SDS-PAGE of LPS^66^. It is possible that AO361_RS10900 functions like *wbpM* in O15 and O17, performing a modification to the OSA of Pse13, and not assembling a core OSA structural component. That specific modification is likely a receptor of Pse05 and Pse04 tailocins, but not Pse06 tailocins.

